# Dynamic, yet well-defined organization of the FUS RGG3 dense phase

**DOI:** 10.1101/2025.07.09.663981

**Authors:** Anton A. Polyansky, Benjamin Frühbauer, Bojan Žagrović

## Abstract

Intrinsically disordered protein regions (IDRs) play a key role in the formation of biomolecular condensates, a ubiquitous mode of cellular compartmentalization, but the underlying microscopic details remain unclear. Here, microsecond-level molecular dynamics simulations and fractal formalism are employed to study at atomistic resolution a model condensate composed of 24 copies of a C-terminal 73-residue arginine- and glycine-rich IDR (RGG3) of fused in sarcoma (FUS) protein. Specifically, RGG3 displays a highly dynamic behavior in the dense phase with only a small configurational entropy loss and a minor slowdown in diffusion as compared to the dilute phase. Despite rapid mixing, short contact residence times and structurally heterogenous binding interfaces in the dense phase, RGG3 exhibits a distinct dynamic binding mode, with statistically defined interaction motifs and a robust multi-scale topology of self-associated protein clusters. The results demonstrate how a well-defined organization of the disordered protein dense phase across scales emerges from highly heterogenous, transient interactions.

## Introduction

Intrinsically disordered regions (IDRs) in proteins play a significant role in different cellular processes, including transcription, translation and signaling^1–5^. The contribution of IDRs to these processes is frequently connected with their involvement in the formation of non-membrane-bound biomolecular condensates^6–8^ such as P-bodies^9^, stress-granules^10^, nucleoli^11^, nuclear bodies^12^ and other^6,13–16^. Importantly, many proteins with large IDRs display well-defined cellular functions, localization, phase behavior and robust binding to multiple targets^17^, while lacking a stable 3D-structural organization. A major open question concerns the emergence of reproducible and robust biological function from the extremely dynamic, disordered behavior of the underlying molecular constituents. For example, it is not clear how a wild-type protein can reversibly form fluid condensates and then, after a single amino-acid substitution in its large IDR, alter its preference in the direction of forming toxic fibrils (see e.g.^18^).

Revision of the structure-function paradigm in the case of disordered proteins brings primary amino-acid sequence to the front. Indeed, a sequence defines how disordered a given region is and, therefore, directly or indirectly impacts all other related functions. Computational approaches for analyzing IDR sequences, including different machine-learning algorithms, have revealed that certain compositional biases, presence of linear motifs, and even a “grammar” can be linked with the level of protein disorderedness^19,20^, interaction preference^21–23^, susceptibility to post-translational modifications^24^, and potential for biomolecular condensate formation^25–27^. However, IDR sequences usually have low complexity and are only weakly evolutionarily conserved, which presents significant challenges to a mechanistic understanding of the sequence-function paradigm. This challenge becomes even harder if one considers the dynamic nature of IDR. Indeed, the folding energy landscape of IDRs is flat and redundant i.e. disordered ensembles consist of many structurally dissimilar, but equally favorable configurations^28^ that exchange rapidly^29^. Thus, sequence properties of IDRs define the nature of their configurational ensembles and understanding of IDR function requires a careful consideration at the ensemble level^17^. This pertains to all levels of organization since, in the case of IDRs, structural heterogeneity and disorder transcend length scales and are equally relevant when it comes to local motifs, complete polymers or polymer networks. Consequently, configurational entropy^28,30,31^ becomes a key parameter that shapes IDR interactions and phase behavior. Specifically, structural reorganization and ordering of IDRs ensembles, which results from binding to a partner^28^ or transitioning into a dense phase, may result in substantial entropic costs^32,33^.

Currently, experimental characterization of the atomistic details of IDRs ensembles and the determinants of their intra- and intermolecular interactions is only possible by using different scattering and NMR techniques, and this only to a limited extent. Experimental limitations can be overcome by the use of computational modeling, specifically molecular dynamics (MD) simulations, which can also be efficiently integrated with the experiment^34^. In particular, MD simulations of IDRs at the all-atom resolution represent a unique biophysical tool since they provide detailed information about the instantaneous position of each atom over realistic, microsecond timescales and, thus, allow direct observation of IDR configurational ensembles and interactions and enable a quantitative assessment of configurational entropy. This has, in particular, been made possible by recent extensive efforts at optimizing the existing MD force fields to generate IDR ensembles in agreement with experimentally derived constraints^35–38^. Thus far, all-atom MD simulations have been used to model conformational behavior of IDRs, interactions between IDRs and other proteins^38,39,40^ or small molecules^41^, and even direct modeling of biomolecular condensate formation^42^ and organization^32,43–45^. All-atom MD simulations, although being restricted in accessible system sizes and timescales as compared to various coarse-grained models widely applied currently in IDR modeling^38^, are much more accurate than the latter in probing the IDR dynamics in different context.

A recent MD study has revealed an intriguingly high level of protein dynamics inside a model condensate consisting of two highly and oppositely charged IDRs^42^. The authors could show that the studied IDRs remain extremely dynamic in the high-density environment (∼300 g/l) of the model condensate, despite a rich network of intermolecular interactions. Moreover, in this mixture, intermolecular contacts were found to be short-lived, with the characteristic relaxation times on the nanosecond scale. In the context of sequence-to-function relationship, are there any well-defined interaction patterns, which can be mapped to the primary sequence and which shape the organization of the highly-dense and dynamic dense phase? The latter aspect is important in order to understand the mechanistic aspects of IDR association and condensate formation. Specifically, sequence patterns of interaction define different relevant parameters such as IDR compactness^46–49^, multivalency^50–53^, and recruitment of RNA and other proteins into condensates^54,55^, while these parameters, in turn, define the mechanism(s) of IDR association and the properties of the ensuing condensates, e.g. their viscoelasticity^56,57^ and topology^32,58^. According to a recent model based on phase separation coupled to percolation, IDRs form a continuum of clusters of different size, whereby at high enough protein concentration clusters grow, merge and coalescence into microscopically visible condensates^32,59^. The distribution of cluster sizes, which are experimentally accessible by scattering techniques such as DLS, can be shifted toward small nanoclusters or large condensates by varying environmental parameters, including protein concentration, temperature or buffer composition^60,61^. Moreover, a newly derived fractal model provides a quantitative link between the atomistic features of an IDRs and the spatial organization of the biomolecular condensate it forms^32^, suggesting that parameters such as condensate topology and density may be directly encoded in IDR sequence.

The RGG3 fragment (residues 454-526, **Figure 1A**) of the human RNA-binding protein (RBP) FUS (fused in sarcoma) is a representative example of an RBP IDR that drives condensate formation and is strongly involved in protein-protein and RNA-protein interactions. FUS is found predominantly in the cell’s nucleus^62^, where it participates in regulating transcription, RNA processing and DNA repair^63,64^. It is also enriched in neuronal cytosol^65^ and cytosolic ribonucleoprotein granules^66^, and it can migrate to the cytosol upon DNA damage as well^67^. The protein is known to form condensates on its own, with both the N-terminal low-complexity (LC) and the C-terminal RGG regions being required for the full effect^68,69^. The RGG-rich region of FUS, in particular, contains a number of charged residues that can drive condensate formation via long-range electrostatic interactions^13^. The abundant Arg and aromatic residues in FUS^70^ have also been suggested to contribute to condensate formation via short-range interactions, as also seen more generally^25,32,48,52,71–73^. Importantly, the C-terminal RGG3 region of FUS is known to form condensates in the presence of sub-stoichiometric amounts of MAPT RNA (1:50) at low salt, with Arg, Tyr and Phe residues being essential for the process^74^. Finally, the FUS RGG3 region includes a small N-terminal patch that is devoid of charges (**Figure 1A**) and the C-terminal NLS, which interacts with different importins. It has been shown that both methylation of Arg residues in RGG3 and importin binding can abolish phase-separation of FUS^75^.

**Figure 1.**
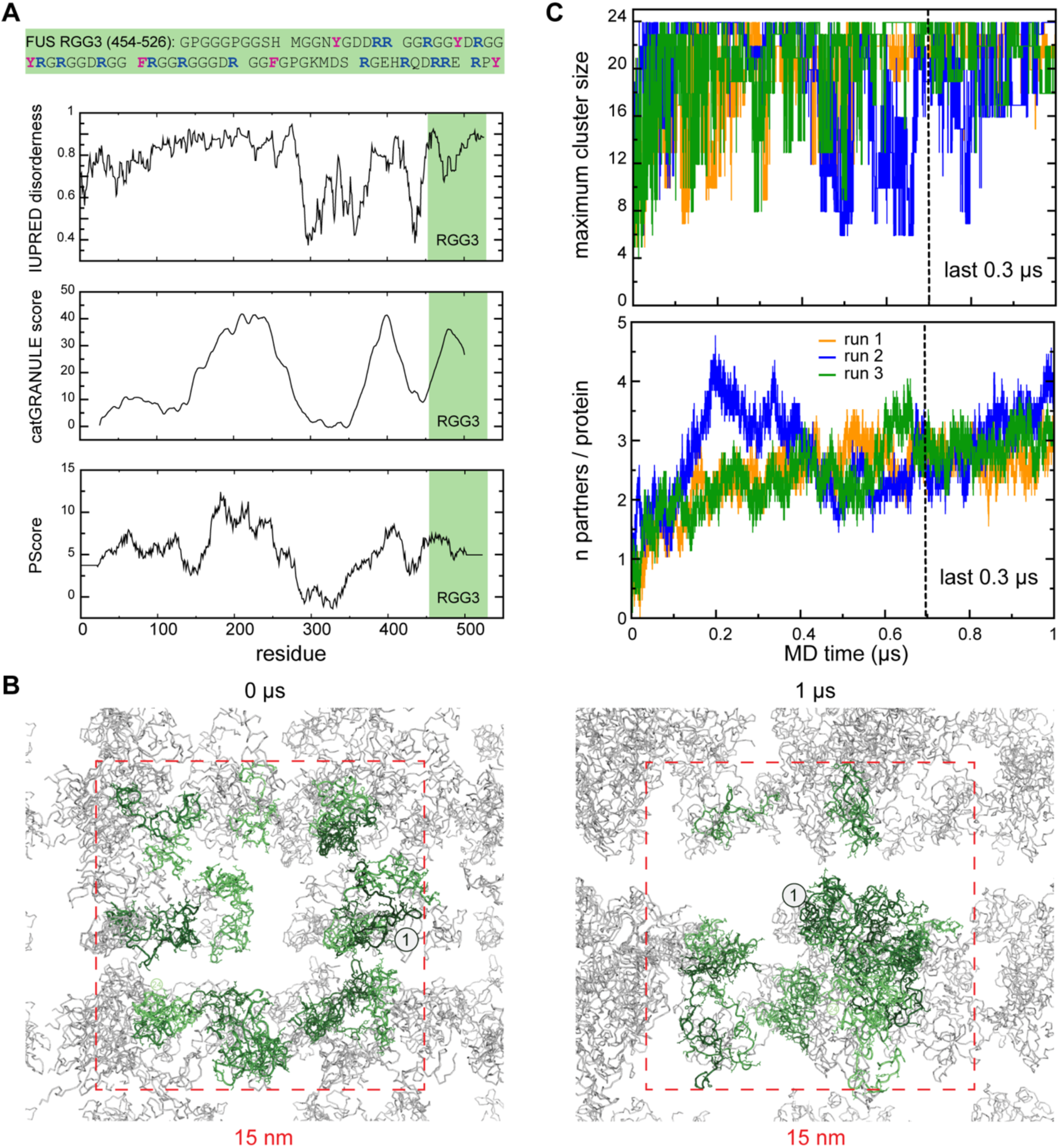
Self-association of the intrinsically disordered RGG3 in the dense phase. **A**. Sequence of RGG3 with aromatic and arginine residues highlighted in magenta and blue, respectively. Panels below show per-residue prediction of RGG3 encoded propensity to intrinsic disorder (IUPRED) and LLPS (catGRANULE score and PScore). **B**. Exemplar snapshots of a multi-copy system (the dense phase) at the beginning (0 µs) and the end of MD simulations (1 µs) with individual protein copies in the central simulation box shown in green and periodic images shown in gray. The simulation box is indicated with a dashed square. Position of a protein copy 1 are indicated. **C**. Time evolution of the largest detected protein cluster size (see Methods for details) and the average valency in the dense phase as calculated for three independent MD replicas of the multi-copy system, which are shown with orange (run1), green (run2), and blue (run3).

To study how a defined and predictable organization of the IDR dense phase can emerge from highly disordered and dynamic interactions, microsecond all-atom MD simulations were employed for systematic investigation of the structure, dynamics, configurational entropy and architecture of intra- and inter-molecular interaction networks of the FUS RGG3 fragment in both a single-copy and 24-copies of the protein in a simulation box, with the latter aimed to model the interior of a biomolecular condensate. The modeling framework used, which enables a multiscale description of the RGG3 condensate, has previously been applied to the disordered Lge1 fragment^32^ and RNA polymerase II C-terminal domain (CTD)^45^.The results demonstrate that RGG3 displays a highly-dynamic and well-mixed behavior with only a small configurational entropy loss upon an isolated protein transitioning to a dense, condensate environment, while at the same time exhibiting a characteristic dynamic self-interaction mode, with statistically defined interaction hotspots and a reproducible topology of self-associated protein clusters across length scales.

## Methods

All MD simulations and analyses were carried out using the GROMACS 5.1.4 package^76,77^, employing Amber99SB-ILDN force field^78^, TIP4P-D water model^35^, and a modeling protocol described elsewhere^32^. Data processing was carried out using custom scripts in GNU bash 4.3.48 and Python 3.6.1. Data visualization and statistical analyses were carried out in R Studio^79^, using the ggplot2 package^80^ for heatmap visualizations and the PK package^81^ for least-squares fitting of a biexponential model to the tail of the pressure tensor autocorrelation functions.

### Single-copy simulations

The starting models for MD simulations of the C-terminal RGG3 (residues 454-526) of human FUS (UniProt ID: P35637) were derived via RaptorX^82^ and Phyre2^83^ webservers. Simulations were set up in tandem for both acquired models using identical parameters. Single molecules were placed in a 9×9×9 nm^3^ box, followed by 1000 steps of energy minimization *in vacuo* under 3D periodic boundary conditions (PBC), alternating conjugate-gradient and steepest descent approaches every 100 steps. The simulation boxes were then filled with ∼24000 water molecules, while Na^+^ and Cl^-^ ions were added to a final concentration of 0.1 M at a net system charge of zero (**Table 1**). Both steps were followed by energy minimization as described above. System equilibration was conducted following a three-step protocol, using one equilibration run in the canonical (NVT) ensemble, with a 0.5 fs time-step for 5000 steps, and two sequential runs in the isothermal-isobaric (NPT) ensemble, with time-steps of 0.5 fs for 25000 steps and 1 fs for and 250000 steps. A leap-frog algorithm was used for integration with PBC. In both energy minimization and equilibration, neighbor-lists were updated every 10 steps, following a Verlet-scheme based grid-search approach. Similarly, in both stages the van der Waals interactions were treated with a twin-range (1.2 nm/1.6 nm) spherical cutoff function, while electrostatics were treated using the Particle-Mesh Ewald method with a real space cut-off of 1.2 nm, 0.12 nm grid and cubic interpolation. The bonds involving H atoms were constrained using LINCS^84^. To control temperature at 310 K, Nose-Hoover thermostat^85^ was used with a relaxation time of 0.5 ps for all steps. To keep isotropic pressure constant at 1.013 bar in the second stage, Berendsen barostat^86^ was used, while Parrinello-Rahman barostat^87^ was used in the final step, with the compressibility of 4.5 x 10^-5^ bar^-^and the relaxation time of 10 ps^1^ in both cases. Coupling was done separately for water and protein molecules in all cases. Two 100 ns-long sampling runs, one for each model (RaptorX and Phyre2), using the same parameters and a 2-fs timestep were set up in order to derive starting structures for multi-copy simulations (see below). Both models converged to a similar radius of gyration (*Rg*) within 30 ns of simulation (data not shown). A 1 µs production run, started using the Phyre2-derived model, was carried out using the same parameters as in the last equilibration run, except for an adjustment of the twin-range spherical cut-off (1.0 nm/1.2 nm) for van der Waals interactions. Finally, three 1 µs MD replicas of the single-copy system were obtained using independent equilibration in each case and a simulation protocol as described above (3 µs in total).

**Table 1.** Details of simulated systems.

| system | box size | MD time, μs | sampling step, ps | proteins | # atoms | | | | |
| --- | --- | --- | --- | --- | --- | --- | --- | --- | --- |
|  |  |  |  |  | protein | water | Na<sup>+</sup> | Cl<sup>-</sup> | total |
| single copy 1 | 9 nm | 1,0* | 10 | 1 | 1010 | 94972 | 44 | 51 | 96077 |
| single copy 1 NVT | 9 nm | 0.05 | 10 | 1 | 1010 | 94972 | 44 | 51 | 96077 |
| single copy 2 | 9 nm | 1.00 | 5 | 1 | 1010 | 95004 | 44 | 51 | 96109 |
| single copy 3 | 9 nm | 1.00 | 5 | 1 | 1010 | 95020 | 44 | 51 | 96109 |
| **24 copies:** |  |  |  |  |  |  |  |  |  |
| run 1 | 15 nm | 1.00 | 2 | 24 | 24240 | 407076 | 203 | 371 | 431890 |
| run 2 | 15 nm | 1.00 | 2 | 24 | 24240 | 406692 | 203 | 371 | 431506 |
| run 3 | 15 nm | 1.00 | 2 | 24 | 24240 | 406692 | 203 | 371 | 431506 |
| run 1 NVT | 15 nm | 0.05 | 10 | 24 | 24240 | 407076 | 203 | 371 | 431890 |
| run 3 NVT | 15 nm | 0.05 | 10 | 24 | 24240 | 406692 | 203 | 371 | 431506 |

\*total simulation time is 1.5 μs , for the final analysis 1 μs fragment used for the consistency

### Multi-copy (dense-phase) system preparation

Three cubic simulation boxes with a side length of 15 nm were each filled with 24 copies of FUS RGG3 by randomly selecting structures from a set of structures drawn from the single-molecule sampling runs at regular intervals, to a final concentration of 11 mM (89.17 mg/ml) in each box. Molecules were placed in a randomized fashion with regards to both molecular identity and orientation on a regularly spaced grid using a custom script ensuring that steric clashes are avoided.

### Dense-phase simulations

Three independent simulations of the 24-copy systems were obtained following the same energy minimization, equilibration and production run protocols as for the single-copy simulations. The simulation boxes were filled with ∼100,000 water molecules, while Na^+^ ions and Cl^-^ ions were added to a final salt concentration of 0.1 M at a net system charge of zero (**Table 1**). Coordinates and energies were written out every 2 ps with the final simulation time of 1 µs per simulation (3 µs in total). Two of the boxes were additionally simulated for 50 ns in the NVT ensemble for viscosity determination. Energies and coordinates were written out every 10 fs and 10 ps, respectively.

### Water simulations

TIP4P-D water was simulated using cubic boxes of different size (edge-lengths of 3, 4 or 5 nm) to determine its *in silico* rheological properties. All boxes were first energy minimized as described above, followed by the addition of Na^+^ and Cl^-^ ions to a final concentration of 0.1 M. After additional energy minimization, equilibration was conducted as for single-copy protein simulations, followed by 100-ns production runs for each system, using the same parameters as for protein NVT simulations and writing out energies and coordinates every 10 fs and 2 ps, respectively.

### In silico rheology

System shear viscosities and translational diffusion coefficients of solute molecules were estimated following the procedures by von Bülow et al.^88^, relying on the previous work of Hess^89^, who explored ways to determine viscosities from simulations, and Yeh & Hummer^90^, who investigated how the system size influences the diffusion coefficients of simulated molecules. Specifically, shear viscosities were extracted from NVT simulations via the Green-Kubo formula:

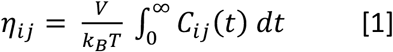

where V denotes the volume of the simulation box and C_ij_ represents the autocorrelation function:

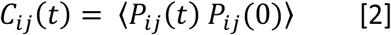

of the pressure tensor elements 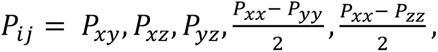, *and* 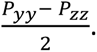

The autocorrelation function was numerically integrated between 0 ps and 1 ps, followed by analytical integration up to infinity. The analytical part of the integral was determined by a least-squares fit of the data between 1 ps and 4 ps to a bi-exponential function:

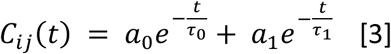

Shear viscosity η was then determined by averaging over the η_ij_ of the evaluated pressure tensor elements.

Translational diffusion coefficients D_t_ were extracted for individual molecules by evaluating the center-of-mass mean-square displacement (MSD) curves, considering that:

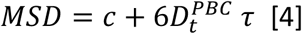

for τ approaching infinity. This equation was fitted in a linear regime between 10 ns and 30 ns for the 24-copy systems, and between 0 ns and 5 ns for the single molecule. The constant c in the formula accounts for an offset caused by non-diffusive behavior over very short time-separations. As established by Yeh and Hummer^90^, the diffusion coefficients extracted from MD simulations are not directly comparable to experimental data, but need to be corrected for size-dependent effects that arise from PBC. Applying this correction, the diffusion coefficient D_t_ can be determined as:

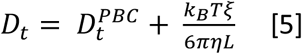

where L is the length of the simulation box, η is the viscosity of the system that the particle is simulated in, and ξ = 2.837297, a term arising from the cubic lattice (see Yeh and Hummer^90^ and references therein). Additionally, to enable a more direct comparison with experimental values, Fennel et al.^91^ proposed to multiply the corrected diffusion coefficient by the ratio of the simulated and experimentally determined viscosities:

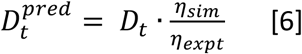

This step should account for inaccuracies in how well the solvent characteristics are reproduced by the water model used. The experimental viscosity at 310 K and 0.1 M salt is η_expt_ = 0.694 mPa*s^91^.

### Trajectory analysis

System energies, *Rgs* and solvent-accessible surface area (SASA) of individual molecules, and the interatomic distances between and within molecules were analyzed using GROMACS^76^. Energy-related terms were analyzed using an output time-step of 2 ps, whereas all other analyses were conducted using a 10 ps step. A distance of 3.5 Å between centers of any two non-hydrogen atoms was used as a cut-off for defining inter-residue contacts. Instantaneous interaction valency of a protein molecule was defined as the number of partners a given molecule was in contact with at a given time point.

### Cluster analysis

A maximum size of protein-protein interaction clusters in the multi-copy simulated system was identified using hierarchical clustering applied to minimum-distance matrices, which were evaluated from each MD trajectory of the multi-copy system sampled at every 10 ps using GROMACS *mindist.* The clustering was done in MATLAB (R2009) using function *cluster* with an applied distance cutoff of 3.5 Å.

### Estimation of expected intermolecular interaction frequencies

The expected frequency of intermolecular interactions as a function of the type of residues involved was determined under the assumption that it depends only on the frequency of the respective residue type in the sequence, according to the following formulas:

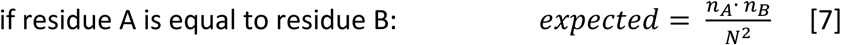

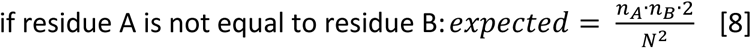

where n_A_ is the number of residues of type A in the sequence, n_B_ is the number of residues of type B in the sequence, and N is the length of the sequence.

### Identification of interaction motifs

Regions with increased contact formation frequency in simulations were determined by assigning a value of interactivity to each residue as the number of contacts (either inter- or intramolecular, depending on the context) of a given residue with all other residues. The interactivity was normalized by the maximum value in a given set for the purposes of visual representation, followed by smoothing of the resulting interactivity-profiles using the Savitzky-Golay filter:

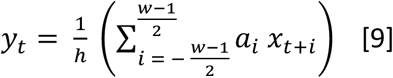

Normalization parameters h and coefficients a_i_ for window sizes w of 5, 7 and 9 were used as suggested by Lohninger^92^. Identification of interaction motifs as the minima in the second derivative of the interaction profiles was also performed via the Savitzky-Golay filter. The minima were selected by rejecting all values that were closer than one standard deviation to the mean of the second numerical derivative of the interaction profile. Shapiro-Wilk test was used to ensure distribution normality. This process was adopted for w = 5, 7, and 9, and each minimum was assigned as the middle point of a putative interaction motif of a window-size w. The final assignment of interaction motifs considered all stretches along the sequence that were found to include a putative interaction motif in at least two different window sizes.

### Error propagation

Error propagation was carried out using the following formula^93^:

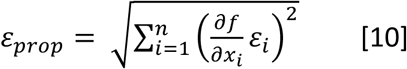

Here, ε_prop_ denotes the propagated error, whereas ε_i_ denotes the error of individual values related by a function f(x_1_, x_2_ … x_n_).

### Entropy calculation

The configurational entropy was estimated using the maximum information spanning tree (MIST) approximation^94^ as implemented in the PARENT program suite^30^ and also described elsewhere^32^. Here, all MD trajectories were converted to Bond-Angle-Torsion (BAT) coordinates. The convergence of configurational entropy (*S*_conf_) was assessed for single-copy systems using cumulative plots with a 50-ns time step. To account for a relatively slow convergence (**Figure S1C**), entropy calculations were carried out for the entire 1-µs trajectories. The relative entropies (Δ*S*_conf_) of a protein in single-copy and multi-copy systems were averaged over all possible combinations of the 3 single-protein copies and the 24 copies in the dense-phase systems. The final values of Δ*S*_conf_ were multiplied by the temperature (T = 310 K) and converted to kcal/mol units.

### Conformational statistics

Conformational dynamics of RGG3 was analyzed using a sliding-window scheme of overlapping pentamer (5-mer) and decamer (10-mer) peptide fragments with single-residue window shifts. For each of the thus defined fragments, all of their local conformations in the multi-copy simulations were combined into a single trajectory (24 µs / 1 ns time step). Pentamer and decamer trajectories were constructed for each multi-copy replica and later combined into a single, master trajectory (72 µs / 3 ns time step).

### Conformational clustering

Conformational clustering on the subsampled pentamer and decamer ensembles was performed using the GROMACS *cluster* utility with backbone RMSD cut-offs of 0.05 nm and 0.1 nm defining the cluster perimeter for pentamers and decamers, respectively. For each peptide fragment, the most populated conformational clusters were identified using combined MD trajectories (also for each independent replica).

### RMSF calculations

Combined pentamer and decamer trajectories were used to calculate backbone atom-positional root-mean-fluctuations (RMSF) for each overlapping peptide fragment using the GROMACS *rmsf* utility. The RMSF value of a middle residue in the analyzed peptide fragments (3^rd^ in pentamers and 5^th^ in decamers) was used to reconstruct RMSF profile for the full-length protein.

### Application of the fractal model

To model and visualize the topology of the self-assembled RGG3 clusters, a previously developed analytical framework assuming the fractal mode of protein self-association^32^ was employed. According to this model, condensate architecture across length scales can be descried as a function of protein valency (*n*) and compactness, or volume fraction (*ϕ*). These two parameters allow a direct assessment of the topological organization of self-assembled protein clusters across scales via the calculation of a fractal dimension (*d_F_*) as follows:

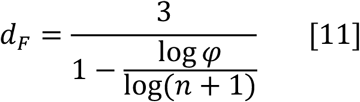

The valency (or the coordination number, *n)* can be directly obtained from the analysis of protein-protein contacts (see above), while compactness (*ϕ)* is defined as the ratio between the molecular (van-der-Waals) volume (*V_mol_*) and the apparent one, as derived from the *Rg*:

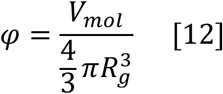

where *V_mol_* is proportional to the molecular weight (*M_w_*) with the used prefactor k as 1.21^32^:

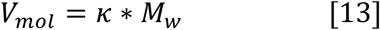

*d_F,,_ Rg,* and *n* were estimated from independent MD trajectories of the multi-copy system and also averaged over the last 0.3 µs. These average parameters were further used to generate a visual representation of the corresponding protein clusters (condensates) using FracVAL algorithm^95^, as described elsewhere^32^. The clusters generated by FracVAL for visualization purposes contained 1024 proteins.

## Results

According to IUPred2A^19^ analysis, (**Figure 1A**), the full FUS RGG3 sequence is predicted to have a high degree of disorder, which is supported by previous experimental evidence^96,97^. Furthermore, RGG3 (**Figure 1A)** exhibits regularly spaced aromatic residues throughout, which are a known characteristic of proteins forming biocondensates^52^, as well as RGG repeats, which are prone to forming π-π interactions, contributing to condensation^26^. Indeed, the compositional bias of RGG3 is reflected in a high predicted propensity towards phase-separation according to catGRANULE^98^ and PSscore^26^ (**Figure 1A**).

### RGG3 self-associates in the dense phase

Upon microsecond MD simulations, a clear tendency towards self-association is observed for RGG3 fragment in the dense phase (**Figure 1B**). Specifically, the highly dynamic individual RGG3 chains form a complex network of intermolecular contacts (**Figure 1B**), characterized by a rapid exchange of binding partners and a formation of protein clusters of different size. The maximum cluster size increases with time in all three replicas (**Figure 1C**) and settles in a relatively wide range (16-24 protein copies) over the last 0.3 µs of simulation time. Interestingly, the system is able to form only a transient percolating cluster, indicating that the contact probability is lower than it is required for a stable single percolating cluster with 24 interconnected copies and robust network transition. Specifically, the probability of proteins to form contacts with neighbors is related to the average valency of interactions, a measure of how many other copies a protein can coordinate. This parameter for RGG3 increases with time in all three multi-copy systems (**Figure 1C**) and stabilizes around 3.3 ± 0.3 over the last 0.3 µs, with a relatively small deviation between the replicas (**Table 2**). Given the convergence in valency over this stretch, the last 0.3 µs of each of the three simulations were used for the analysis of other equilibrium properties of RGG3 throughout this work.

**Table 2.**
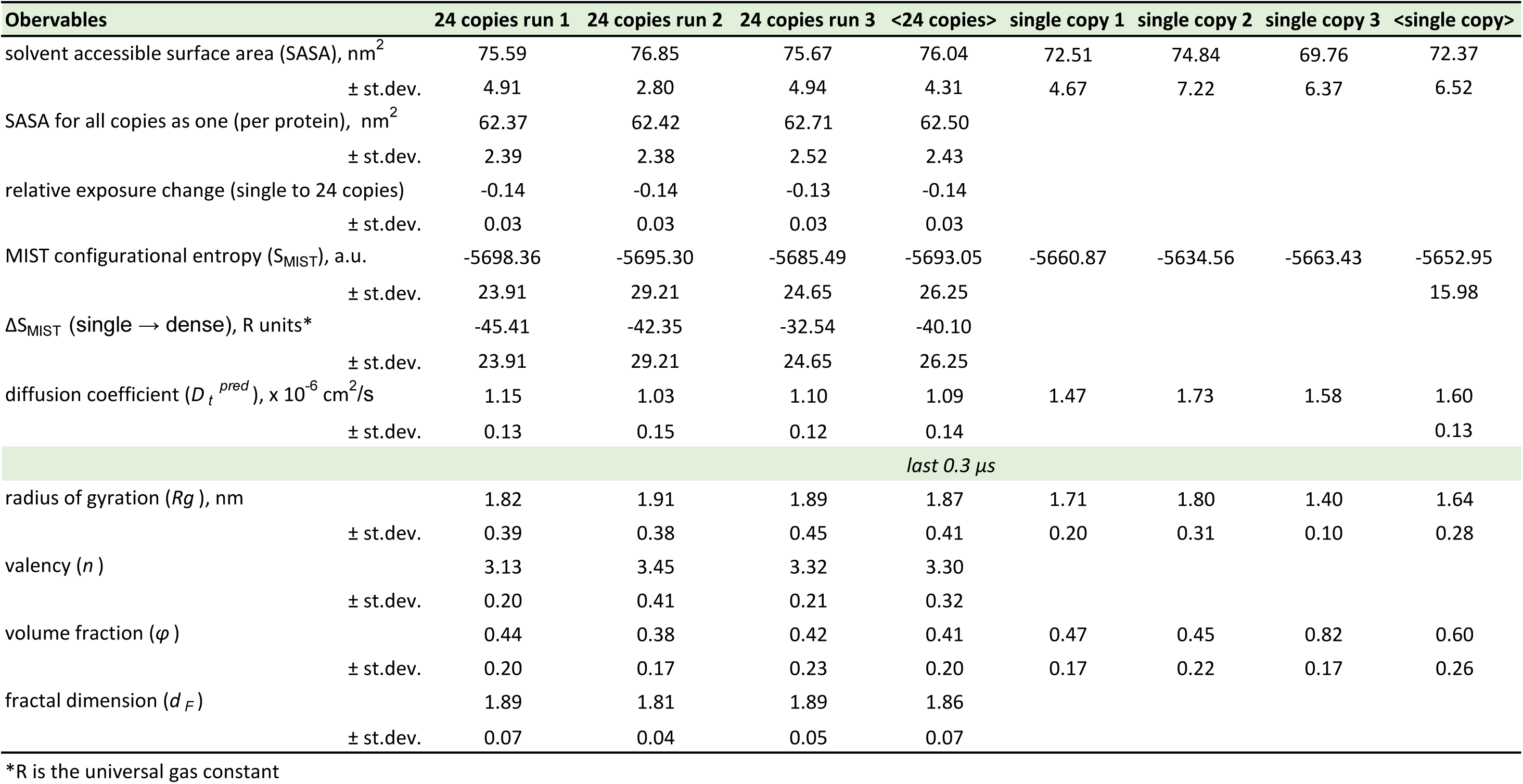
Summary table on observables derived from analysis of MD trajectories.

| Obervables | 24 copies run 1 | 24 copies run 2 | 24 copies run 3 | <24 copies> | single copy 1 | single copy 2 | single copy 3 | <single copy> |
| --- | --- | --- | --- | --- | --- | --- | --- | --- |
| solvent accessible surface area (SASA), nm<sup>2</sup> | 75.59 | 76.85 | 75.67 | 76.04 | 72.51 | 74.84 | 69.76 | 72.37 |
| ± st.dev. | 4.91 | 2.80 | 4.94 | 4.31 | 4.67 | 7.22 | 6.37 | 6.52 |
| SASA for all copies as one (per protein), nm<sup>2</sup> | 62.37 | 62.42 | 62.71 | 62.50 |  |  |  |  |
| ± st.dev. | 2.39 | 2.38 | 2.52 | 2.43 |  |  |  |  |
| relative exposure change (single to 24 copies) | -0.14 | -0.14 | -0.13 | -0.14 |  |  |  |  |
| ± st.dev. | 0.03 | 0.03 | 0.03 | 0.03 |  |  |  |  |
| MIST configurational entropy (S<sub>MIST</sub>), a.u. | -5698.36 | -5695.30 | -5685.49 | -5693.05 | -5660.87 | -5634.56 | -5663.43 | -5652.95 |
| ± st.dev. | 23.91 | 29.21 | 24.65 | 26.25 |  |  |  | 15.98 |
| ΔS<sub>MIST</sub> (single → dense), R units* | -45.41 | -42.35 | -32.54 | -40.10 |  |  |  |  |
| ± st.dev. | 23.91 | 29.21 | 24.65 | 26.25 |  |  |  |  |
| diffusion coefficient ($D_t^{pred}$), x 10<sup>-6</sup> cm<sup>2</sup>/s | 1.15 | 1.03 | 1.10 | 1.09 | 1.47 | 1.73 | 1.58 | 1.60 |
| ± st.dev. | 0.13 | 0.15 | 0.12 | 0.14 |  |  |  | 0.13 |
| *last 0.3 μs* |  |  |  |  |  |  |  |  |
| radius of gyration ($R_g$), nm | 1.82 | 1.91 | 1.89 | 1.87 | 1.71 | 1.80 | 1.40 | 1.64 |
| ± st.dev. | 0.39 | 0.38 | 0.45 | 0.41 | 0.20 | 0.31 | 0.10 | 0.28 |
| valency ($n$) | 3.13 | 3.45 | 3.32 | 3.30 | | | | |
| ± st.dev. | 0.20 | 0.41 | 0.21 | 0.32 |  |  |  |  |
| volume fraction ($\varphi$) | 0.44 | 0.38 | 0.42 | 0.41 | 0.47 | 0.45 | 0.82 | 0.60 |
| ± st.dev. | 0.20 | 0.17 | 0.23 | 0.20 | 0.17 | 0.22 | 0.17 | 0.26 |
| fractal dimension ($d_F$) | 1.89 | 1.81 | 1.89 | 1.86 | | | | |
| ± st.dev. | 0.07 | 0.04 | 0.05 | 0.07 |  |  |  |  |

\*R is the universal gas constant

### RGG3 dense phase is dynamic and structurally heterogeneous

The instantaneous valency of RGG3 ranges between 0 and 9 binding partners per protein in all three multi-copy simulations (**Figure S1A**), reflecting a variety of bound configurations adopted by the protein in a crowded environment. Importantly, such interactions are characterized by a frequent partner exchange, which is well illustrated by the number of possible intermolecular pairs realized as function of time (**Figure 2A**). In particular, over 90% of all possible pairs of individual RGG3 chains establish at least one direct contact in the course of 1-μs multi-copy simulations, while approximately 60% of all possible pairs contact each other at least once over the last 0.3 µs. The interaction dynamics can also be captured by the rate of exchange of binding partners via valency autocorrelation function. Specifically, in all three multi-copy simulations, valency persists at a given level between 30 and 40 ns on average, with any memory of valency dissipating by ∼150 ns (**Figure 2B**). In fact, although the average valency is only ∼3.3, each protein chain interacts directly with ∼14 different partners over the last 0.3 µs, whereby contacts with more than 50% of partners last on average shorter than 1 ns each (**Figure 2C**). Clearly, contacts between different RGG3 copies form and dissolve rapidly and there is a high degree of mixing on the time scale of our simulations, creating a foundation for a detailed statistical analysis of interaction motifs (see below).

**Figure 2.**
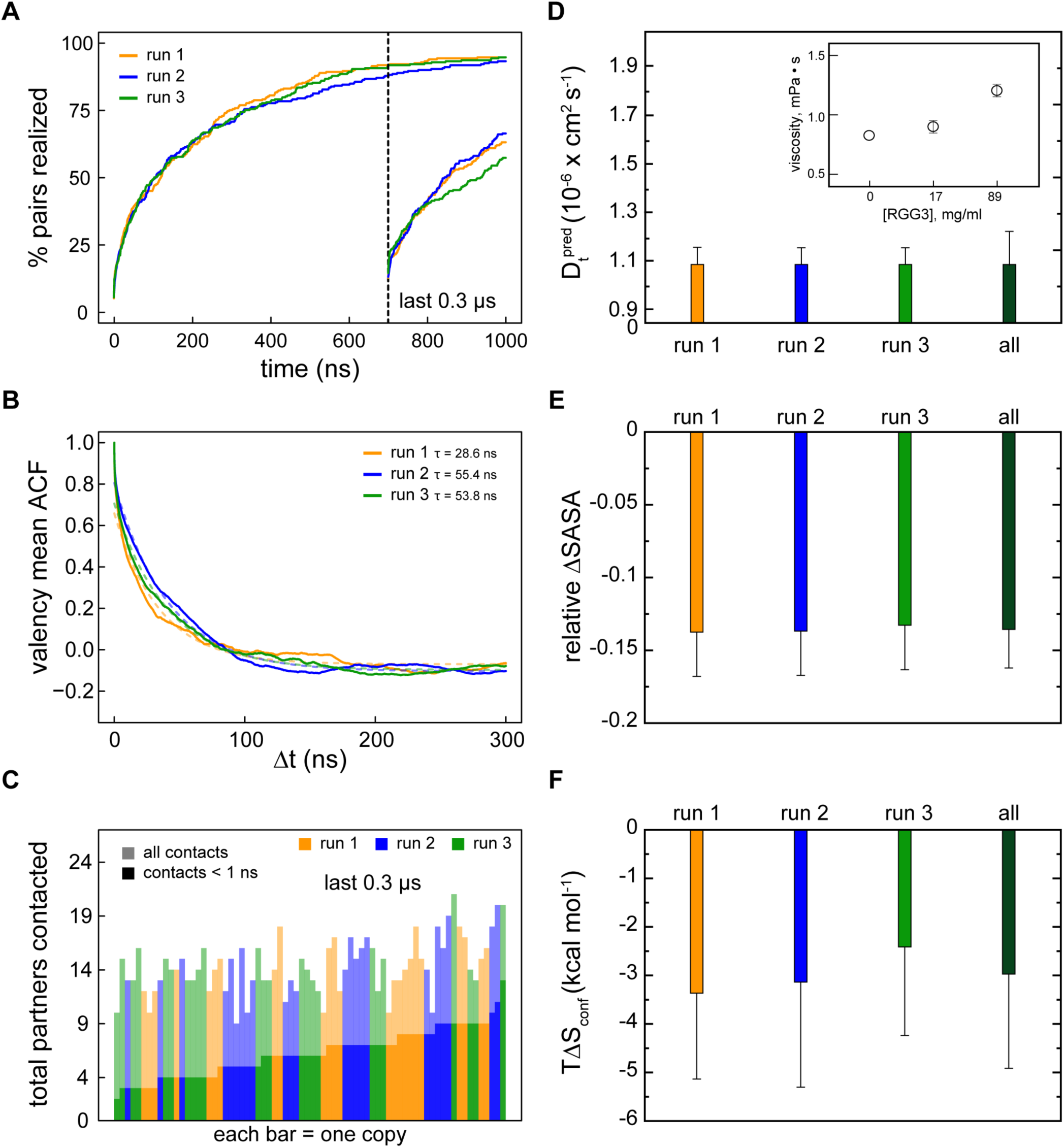
Dynamic and structurally heterogeneous nature of the RGG3 dense phase. **A**. Fraction of all 276 (=24*24/2) possible pairwise intermolecular contacts that were established at least once after t=0 µs or t=0.7 µs in the three multi-copy system replicas. **B**. Average autocorrelation function (ACF) of protein valencies as a function of time separation Δt along each independent MD trajectory. Single exponential fits of ACF curves are shown with dashed curves. Corresponding characteristic times of the fits are indicated in the legend. **C**. Number of binding partners for each the 72 individual molecule over the last 0.3 µs of multi-copy simulations, with the contacts longer than 1 ns shown in darker colors. **D**. MD-derived translational diffusion coefficients (see Methods for details) of RGG3 in the dense phase. The value was averaged between all 24 protein copies in the system. Error bars depict standard deviations. *Inset:* Shear viscosity values obtained from the analysis of the pressure tensor autocorrelation functions (see Methods for details) and shown as function of RGG3 mass concertation. The average values over different pressure tensor elements are shown together. Error bars depict standard deviations. **E.** Average relative changes in the solvent accessible area (SASA) of RGG3 in isolation upon transitioning into the dense phase. Averaging was done for SASA differences in all possible combinations between three independent runs of single RGG3 and 24 protein copies in each complete 1µs MD trajectory of the dense phase. Error bars depict standard deviations. **F**. Average changes in the configurational entropy (Δ*S_conf_*) of RGG3 in isolation upon transitioning into the dense phase. Entropy values are given in energy units (*T*Δ*S_conf_*, = 310 K) and were obtained using complete 1 µs MD trajectories. Averaging was done for entropy differences in all possible combinations between three independent runs of single RGG3 and 24 protein copies in the in each MD replica of the dense phase. Error bars depict standard deviations.

Highly dynamic interactions with frequent, nanosecond-level binding and unbinding suggest that RGG3 may diffuse rapidly in the crowded environment. A linear dependence of protein MSD on time, indicative of an ideal diffusive behavior, was identified between 10-30 ns and used for deriving diffusion constants in the multi-copy systems (**Figure S1B**). Interestingly, the single-copy simulations exhibit no significant linear regions in the MSD vs. time curve. Fitting the MSD-curves in the range from 0-5 ns, as performed before^88^, still allowed estimation of the translational diffusion constant in those cases. Overall, both viscosity and diffusion constants derived from simulations (**Table 2**, **Figure 2D**) are in good agreement with the behavior expected from previous experimental and theoretical results. Specifically, our water model exhibits an apparent viscosity of 0.83 ± 0.01 mPa*s, close to an experimental estimate of 0.69 mPa*s^91^ (**Figure 2D**, **Table S1**), with the value increasing by ∼10% and 50% for single-copy and multi-copy RGG3 systems, respectively. Furthermore, the translational diffusion constants of proteins in our simulations agree with previous simulation results^32,88^. Interestingly, the average diffusion constant of RGG3 in the single-molecule context (1.60 ± 0.13 x 10^-6^ cm^2^/s) reduces by only 32% in the dense phase (1.09 ± 0.14 x 10^-6^ cm^2^/s) (**Table 2**). Thus, transient interactions with other protein partners and association into clusters slow down the diffusion of RGG3 only marginally.

Formation of the protein network results also in a 14% reduction of protein SASA in the dense phase as compared to the single-copy context (**Table 2**). Specifically, the average SASA for RGG3 over all three multi-copy system is 62.5 ± 2.4 nm^2^, with very small differences between the replicas, while this variation is a few times higher for single-copy replicas (**Figure 2E**). The systematic reduction of SASA upon crowding is also reflected in the thermodynamics of RGG3 self-association. Thus, calculations show that the configurational entropy of disordered proteins systematically and reproducibly decreases upon crowding, with an average *TΔS_conf_* per molecule of -3.0 ± 2.2 kcal/mol (**Figure 2F**). However, due to the very dynamic organization of the protein network, this effect is significantly smaller as compared to the Lge1 (1-80) fragment (*TΔS_conf_* = -6.6 ± 3.5kcal/mol), which was shown to robustly form condensates *in vitro* and stable percolating protein clusters in MD^32^. Interestingly, the reduction of the RGG3 configurational entropy upon crowding in all three multi-copy systems is linearly proportional to the relative reduction of its SASA (**Figure S1D**). Since the decrease in the RGG3 solvent-exposed surface is related to the release of water molecules from it, it is likely that solvent entropy increases proportionally to the reduction of protein configurational entropy and that this may contribute to the driving force of condensate formation in a manner that is analogous to the hydrophobic effect in protein folding.

### Interaction motifs in RGG3 can be statistically defined

To study which regions in RGG3 are involved in the above interactions, a detailed analysis of all pairwise intermolecular contacts was performed (**Figure 3A**). For better statistics, the averaging interval of MD trajectories was extended to 0.4 µs in the contacts analysis. Consistent with their high dynamics, RGG3 chains interact with each other throughout the entire sequence, but to a variable degree. To allow identification of these regions, the raw profile of contact formation frequencies as a function of sequence position (**Figure 3B**) was processed via the Savitzky-Golay filter (**Figure 3C, Figure S2B, C**). Specifically, the N-terminus of RGG3 (residues 1-19) exhibits pronounced interactivity in all three replicas, with several additional regions also featuring. Remarkably, despite high structural heterogeneity of simulated systems, an alignment of individual interactivity profiles from different replicas (**Figure 3D**, top) results in a good overall agreement, with pairwise Pearson Rs around 0.8 or greater. This is indicative of a strong convergence when it comes interaction propensities. An exception can be seen in the third peak, which is located around residue 45 in run 1, while being shifted noticeably to the left in runs 2 and 3. Notably, a sequence interaction profile with the cleanest separation between interaction hotspots is obtained if one focuses just on conformers with valency=1 i.e. chains that interact with one other partner only at a given time point (**Figure 3D**, bottom). These conformers may inform on the residues involved in the early contact formation. Finally, one can use the Savitzky-Golay filter to identify sequence regions that mediate the formation of intermolecular contacts in an unbiased way. Using this approach, a repeating DRGG(F/Y) motif was identified within the RGG-rich region as a driver of intermolecular contact formation in RGG3 (**Figure 3E, Figure S2D**). Interestingly, the DRGGF motif closest to the N-terminus is determined with the highest confidence, while the subsequent motifs show a continuously decreasing interactivity. A possible explanation for this can be found in the residues surrounding each motif: the first is flanked by the highly interactive Tyr and Arg, while the third motif is surrounded by Gly residues, which likely cannot stabilize contacts to the same degree.

**Figure 3.**
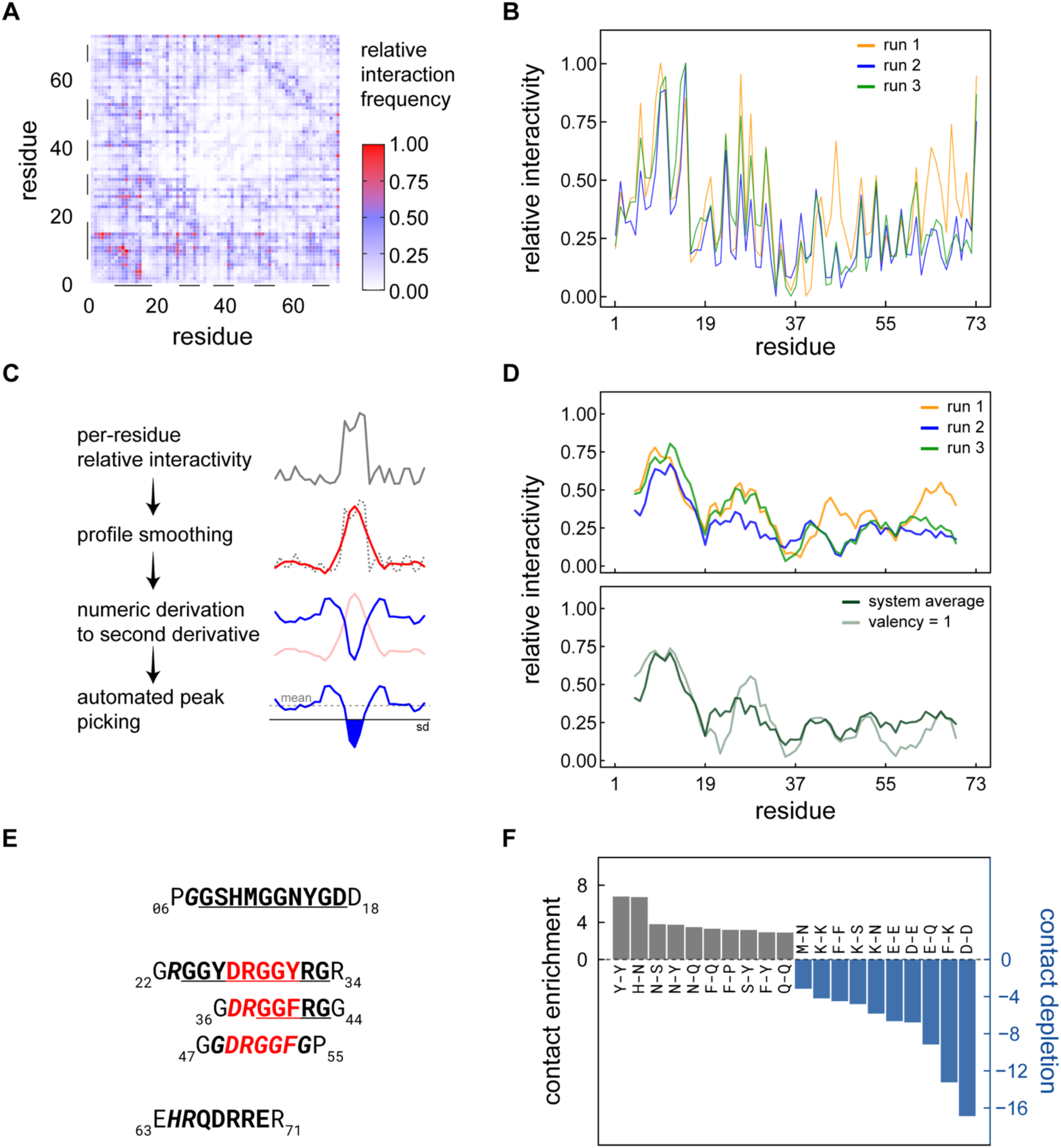
Statistically defined interaction motifs in RGG3. **A.** Per-residue heatmap of intermolecular interactions. Bars highlight positions that are subsequently identified as significant drivers of intermolecular interactivity. **B.** Unfiltered (raw) relative residue interactivities (contact formation frequencies) of RGG3 in the dense phase. **C.** Overview of Savitzky-Golay filtering and peak-picking (see Methods for details). **D.** Contact frequencies profiles of RGG3 for individual multi-copy simulation replicas and comparison of interactivity profiles at all valencies and at valency 1. **E.** Alignment of interacting regions in RGG3 as identified by Savitzky-Golay filtering: the common DRGG(F/Y) motif was identified in all three interaction “hotspots” located in the RGG-rich region of the protein. **F.** The ten most enriched and depleted types of pairwise intermolecular contacts involving molecules with a valency of 1 in the RGG3 dense phase.

The 20 most frequent types of residue-residue contacts between different copies of RGG3 are shown in **Figures S2E**. Many of these involve residues that are abundant in the protein’s sequence, most notably Gly. To account for this, the 10 most enriched and the 10 most depleted pairs of residues among the observed contacts were identified and normalized by their abundance in the RGG3 sequence (**Figure 3F**, see **Figure S2F** for the statistics without filtering by the valency). The contacts involving Tyr are significantly enriched in the multi-copy system, together with the contacts that are stabilized by hydrogen bonding, such as those involving Asn. Depleted contacts, on the other hand, generally tend to involve polar and charged residues. Notably, the depletion of Phe-Lys interactions is consistent with previous findings in other proteins^99^. Finally, both the enriched and the depleted contacts include residues of different size. This is important as amino acids with larger side chains would be more likely to contact each other by accident in a crowded environment. Thus, the relationship between residue size in terms of surface area and per-residue intermolecular interactivity were analyzed and compared for the whole system and for contacts at valency=1 only (**Figure S2A**). The correlation between residue size and interactivity in the full system (r = 0.55) appears to be guided by the residues on the two ends of the size spectrum, and shows outliers among the medium-sized residues. At valency 1, however, the correlation largely disappears (r = 0.25), suggesting that the interactions are established more specifically in that case.

### RGG3 interaction motifs are intrinsically disordered

In line with the predicted disorder of RGG3, structural clustering of the protein as a whole showed a marked absence of any conformational preference (data not shown). However, it is possible that the interaction regions adopt locally well-defined conformations upon binding. Therefore, RMSD-based structural clustering was performed on consecutive 10-residue fragments along the RGG3 sequence (see Methods) to identify the most populated conformational cluster for each (**Figure 4A**). The most highly occupied clusters i.e. the most structurally well-defined regions, are located at the protein’s C-terminus, with an occupancy rate approaching 20%. However, the top conformational clusters that contain the identified N-terminal interaction hotspots or DRGG(F/Y) motifs exhibit mostly below-average occupancies, and a clear conformational similarity between the top clusters of the motifs is absent. Curiously, the low interacting regions (“spacers”) appear to be conformationally better defined i.e. exhibit increased occupancies of the top conformational clusters (**Figure 4A**). The total number of clusters in individual 10-mer stretches was also analyzed (**Figure 4B**). The high number of clusters at the N-terminus and the low number of clusters at the C-terminus are consistent with the respectively low and high occupancies of the top clusters at these positions. The interaction motifs also overlap with the positions of highly fluctuating sequence regions (RMSF peaks, **Figure 4C**), especially for the first two peaks. Interestingly, RMSF profiles generated for the combined MD trajectories (see Methods) represent a characteristic dynamic fingerprint of RGG3 in the dense phase, where position of the regions with high conformational heterogeneity are rather independent from the size of the applied sliding window (not shown). Altogether, the systematically high number of local conformational clusters along the entire RGG3 sequence suggests that binding events that involve both high- and low interacting regions are structurally heterogeneous and devoid of well-defined configuration, reminiscent of what has been termed a “fuzzy complex”^100^.

**Figure 4.**
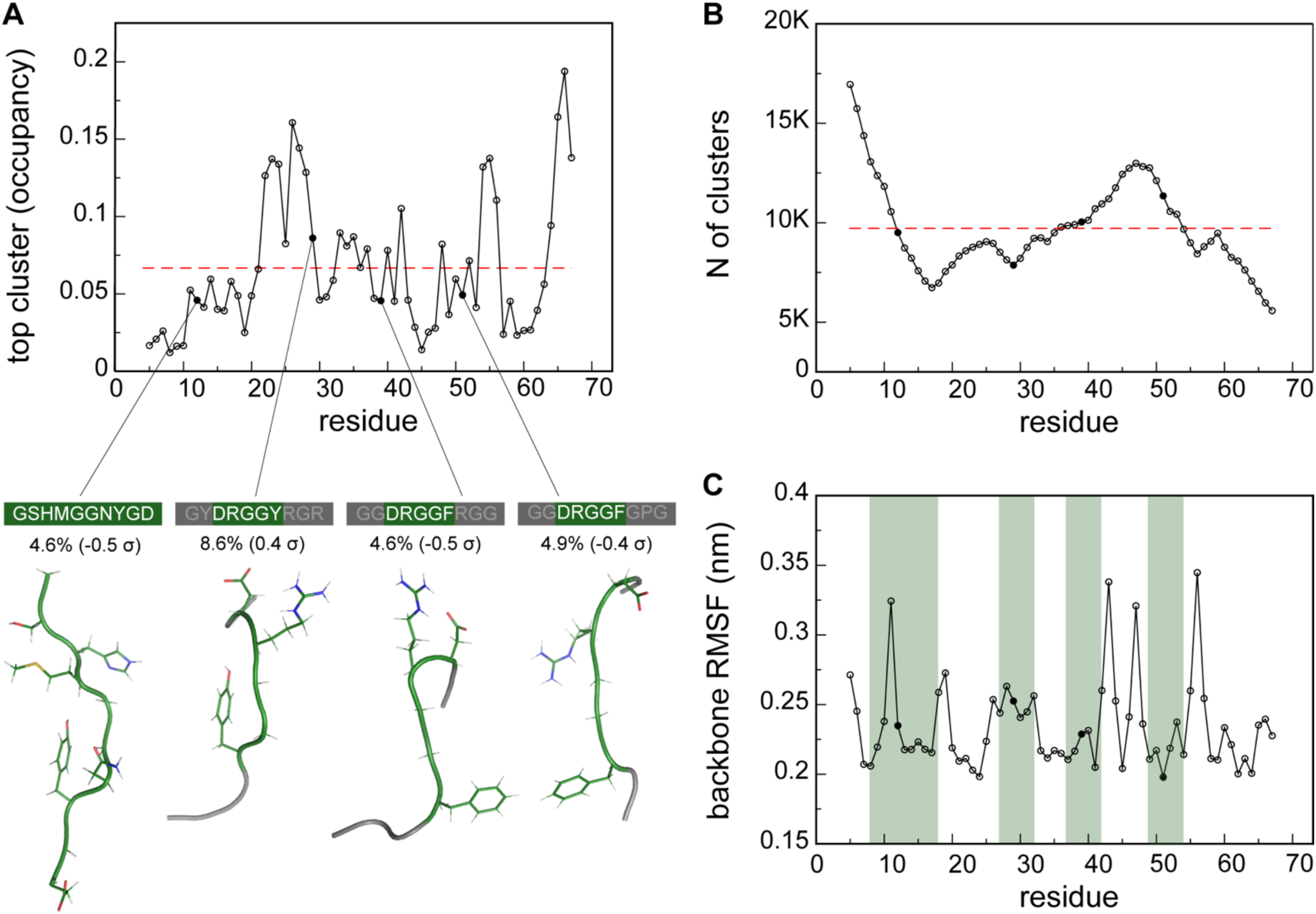
Disordered nature of RGG3 interaction motifs. **A.** Relative occupancy of the top conformational clusters of 10-residue fragments in RGG3. The average occupancy of top clusters is indicated with a dashed line, while central structures of top conformational clusters for the identified interaction motifs are shown below. Relative occupancies for the top clusters of the interaction motifs (in %) and deviation from the baseline (in standard deviations) are given below the motif sequences. For short interaction motifs, sequence neighbors within 10-mer fragments are shown with gray color. **B.** Total numbers of delineated conformational clusters for each 10-mer fragment. The average is indicated with a dashed line. **C.** Average per-residue root-mean-squared fluctuations (RMSF), using a sliding window of 10 residues. Positions of Interaction motifs are highlighted in green.

### RGG3 clusters exhibit robust scaling

To probe the topological organization of RGG3 protein clusters across different length scales and translate the observations obtained for the relatively small simulated systems to the sub-micrometer scale, the previously developed fractal model^32^ was employed (see Methods). According to the model, protein compactness and interaction valency define the spatial organization, or “dimensionality” of the self-associating clusters, across different length scales. Specifically, the fractal dimension (*d_F_*) of such protein clusters can be directly calculated from the values of the interaction valency (*n*) and the corresponding compactness of a protein (*ϕ*) using Eq. 11 (see Methods). Overall, RGG3 in the single-copy simulations samples relatively compact conformations, with an average <*Rg>* of 1.64 ± 0.28 nm for all replicas over the last 0.3 µs of MD (**Figure S3A, Table 2**). This is ∼30% smaller than what would be expected for a random-coil peptide of this size according to a simple scaling law^101^. The observed compactness of the single-copy RGG3 indicates that intramolecular interactions shape the configuration of the protein to a significant degree. This value increases by ∼14% in the crowded environment to 1.87 ± 0.41 nm, as averaged over all copies in all replicas (**Figure S3B, Table 2**). Thus, the dense phase promotes spatial extension of the RGG3 conformations due to intermolecular protein-protein interactions and intercalation in the emerging protein network. Interestingly, the average *Rg* in the multi-copy system displays a stable behavior along the most part of MD trajectories in all three replicas (**Figure 5A**), in contrast to the single-copy simulations where fluctuations in *Rg* value are much higher (**Figure S3C**). Thus, the compactness of RGG3 is robustly defined in the crowded environment. This is reflected in the fact that the fractal dimension *d_F_* exhibits a well-defined, nearly identical value for the three multi-copy replicas (**Figure 5B**), with an average value of 1.86 ± 0.07 over the last 0.3 µs (**Table 2**). Therefore, while being very dynamic, the RGG3 protein network is expected to exhibit a very stable topological organization across scales, with the fractal dimension below 2. This is consistent with the network exhibiting a low-dimensionality with a relatively loose organization. Such architecture, as predicted by the fractal model, should be valid for clusters of any size: indeed, the scaling between cluster *Mw* and its *Rg* remains constant and is very similar between all three multi-copy systems (**Figure 5C**). Finally, reconstruction of the 3D organization for a cluster of 1024 RGG3 proteins with Mw of 7.7 MDa clearly demonstrates the relatively loose organization of the protein network (**Figure 5D**) and a very similar large-scale architecture for the three independent MD replicas of the dense-phase system (**Figure S3D**).

**Figure 5.**
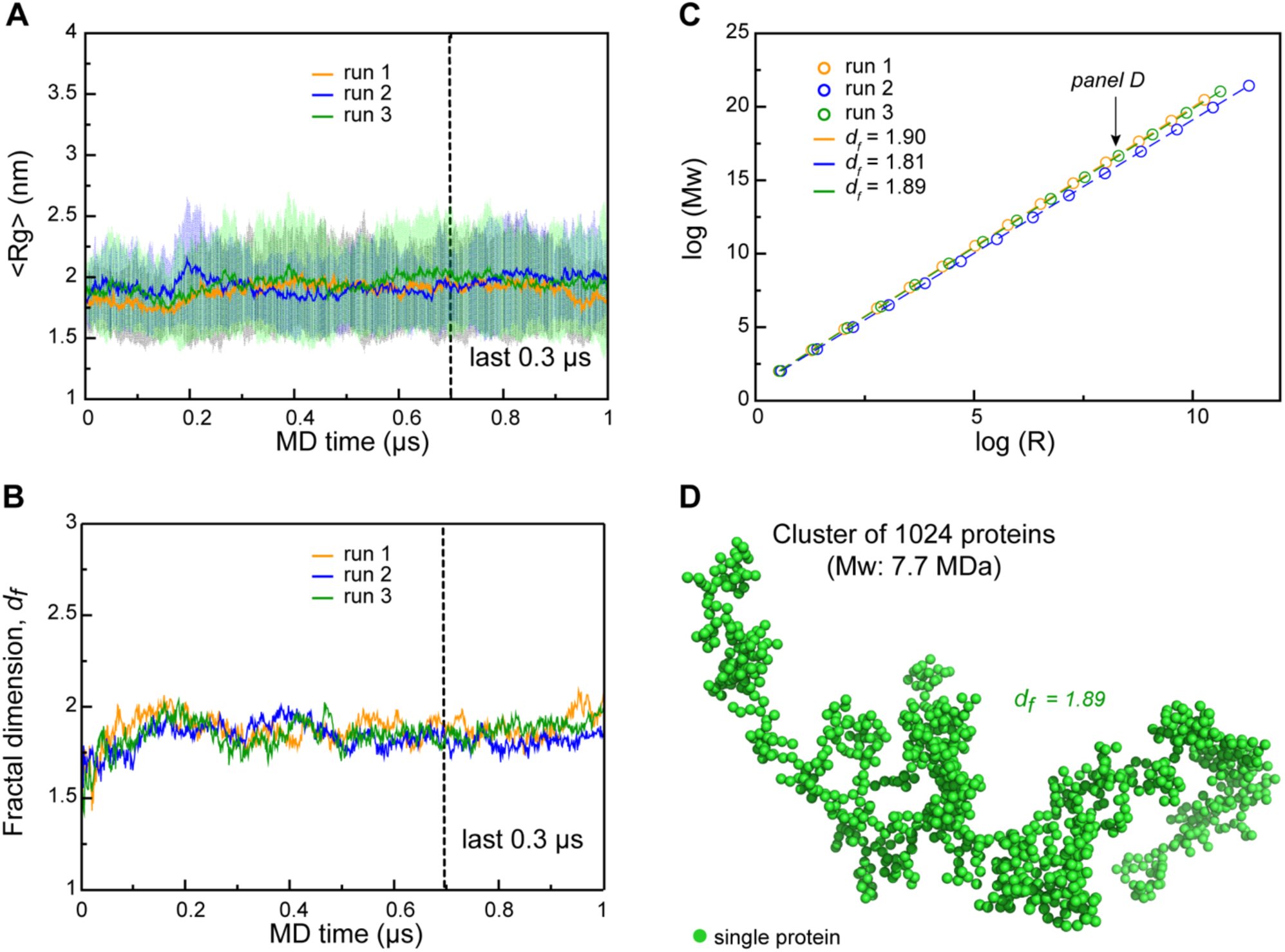
Robust scaling of RGG3 clusters. **A.** Time evolution of the average radius of gyration (*Rg*) in the RGG3 dense phase. The average values over all 24 copies are shown together with the standard errors of the mean. **B**. Time evolution of the fractal dimension (*d_F_*) in the RGG3 dense phase. The fractal dimension is calculated using average *Rg* and valency values (eq. 11-13, see Methods) between 24 protein copies at each time step of MD trajectories. **C.** Power-law dependence between mass and size of RGG3 clusters at different iterations of the fractal model (see Methods for details) with the applied valency and compactness corresponding to their average values over the 24 simulated protein copies and the last 0.3 µs of each MD replica of the dense phase. Dashed lines show linear regression for the log *R* vs. log *Mw* plot with the corresponding *d_F_* values indicated in the legend. **D.** A representative coarse-grained 1024 particle cluster obtained by FracVAL algorithm (see Methods). The cluster was reconstructed using the *d_F_* and the averaged *Rg* value at the last 0.3 µs of the dense phase MD simulations (run2).

## Discussion

A detailed characterization of the structure, dynamics and interactions of biomolecular condensates at an atomistic scale is an open challenge of both fundamental and applied significance. Here, systematic all-atom MD simulations were employed to provide a detailed view of the dense phase of a biologically relevant IDR fragment, gaining insight into what the driving forces behind phase-separation involving IDRs may be. The simulated RGG3 system is characterized by rapid mixing and a dynamic exchange of intermolecular contacts, with individual proteins remaining structurally disordered throughout simulations. Remarkably, however, the intermolecular contacts in the RGG3 dense phase exhibit robust, statistically well-defined features. Specifically, it was shown that a repeating motif, DRGG(F/Y) drives the formation of intermolecular contacts between different copies of RGG3 via interactions with the highly interactive N-terminal region i.e. a “sticker”^8,52^ . It is likely that the initial formation of contacts is guided by Arg residues, while the aromatic residues provide stability to the dynamic network of interactions.

Analysis of the present MD simulations using the fractal model of condensate architecture across scale shows that the simulated RGG3 system exhibits a remarkably robust and predictable behavior, with the three independent simulations of the dense phase resulting in almost identical values of the fractal dimension *d_F_* and, consequently, almost identical scaling behavior (**Figures 5C**, **5D**, **S3D**). This entirely reflects the fact that the interaction valency and the compactness of individual RGG3 conformers in the dense phase exhibit well-converged, similar values across the different, independent simulations. More generally, this also illustrates a major feature and advantage of the fractal organization of the protein dense phase: in such organization, the large-scale architecture of the complete network is a direct consequence of the intrinsic features of individual chains it consists of i.e. their valency and compactness, which in turn are encodable in protein sequence. Finally, the fractal nature of the protein dense phase implies formation of clusters at different length scales, in agreement with the observed heterogeneous distribution of clusters in subsaturated solutions of FUS-EWSR1-TAF15 family of proteins^102^.

Recently, it was proposed that the regular spacing of aromatic stickers in phase-separating IDPs weakens the strong interaction between aromatics through the favorable solvation of the intervening spacers and in this way optimizes the balance between desirable phase separation and undesirable aggregation^8,52^. The present results suggest that, additionally, polar residues participate significantly in contact formation and that the free-energetic difference between contacts involving sticker regions and those involving spacer regions may only be minor. Thus, the contacts involving sticker regions make up just over half of all intermolecular contacts (**Table S2**), while making up 40% of the sequence. Conversely, spacer-spacer contacts account for almost 47% of all interactions. This provides a novel counterpoint to the canonical sticker/spacer picture, in which spacer interactions are strongly disfavored. Rather, it is likely that all parts of a typical IDR participate in interaction, but with a continuum of interaction strengths and residence times. However, it is also possible that the present simulations capture primarily the early events in condensate formation and that the interactions between stickers get stronger as condensates mature. This caveat also pertains to all structural and dynamical features of condensates analyzed herein.

A number of mutations that are commonly identified in FUS-related pathologies cluster in its C-terminally located NLS and are mapped on the sequence in **Figure 1A**. Interestingly, the NLS contains two motifs identified by the analysis, _49_DRGGF_53_ and _66_QDRRE_70_. What is more, pathological mutations in these regions or their immediately flanking residues (G54D, R68C, R68G, R68H or R69G) invariably involve perturbation of charge states. It is possible that these mutations affect intermolecular contact formation in FUS condensates by modulating electrostatic interactions and exert their deleterious effect in this way. Future mutagenesis work could provide a test of this hypothesis.

Here it is important to emphasize that RGG3 was shown to phase separate in the presence of sub-stoichiometric amounts of RNA (1:50) only and, with nucleotides or RNA absent, our simulations may not provide a complete picture of how RGG3’s residues drive phase separation. However, there are multiple lines of evidence suggesting that protein-protein interactions, as studied herein, play a significant role in the process, including: 1) the high intrinsic propensity of the RGG3 sequence towards phase separation as captured by different algorithms (**Figure 1A**), 2) the central role of the fragment in the phase separation of the complete FUS, with or without RNA^74^.

An accurate analysis of IDRs via MD simulations poses significant challenges when it comes to both the force fields and the degree of sampling required. The choice of the Amber99SB-ILDN force field^78^ in combination with the TIP4P-D water model^36^ ensures that the simulated trajectories capture well the dynamic and disordered nature of RGG3. On the other hand, some protein-protein interactions are still difficult to capture even by the latest force fields^36^, including π−π interactions^26^, which are thought to be a key driver of IDR phase separation. In particular, the strong enrichment of aromatic interactions seen in RGG3, may to some extent be due to a force-field bias, which could be magnified in a crowded environment, especially at high valency. On the other hand, our simulations also clearly point to polar residues as being involved in establishing early, specific contacts between individual RGG3 molecules. Such interactions may guide molecules to bind to specific sites, while a different subset of residues is responsible for contact stabilization.

The second challenge in simulating condensates is posed by a high degree of sampling needed to adequately capture equilibrium behavior. The decision to assign almost 2 µs of MD simulation time to equilibration and combine three independent trajectories into a joint ensemble was taken precisely in order to capture the true equilibrium behavior of the RGG3-PY condensate. Importantly, in each of the trajectories, RGG3-PY was simulated in multiple copies, greatly improving the degree of conformational sampling: altogether, we simulated a total of 72 independent copies of the protein, reaching a combined total of ∼30 µs of sampling in the equilibrated state. Additionally, rapid partner exchange and transient formation of intermolecular contacts have enabled a statistically well-converged characterization of relative interaction propensities. Finally, we observed a close agreement of the three independent trajectories in terms of the average number of binding partners per protein and, most importantly, residue-based interactivity profiles. While it is not possible to discount a qualitatively different behavior of the condensate on a time scale beyond microseconds, such convergence strongly suggests that the obtained data accurately represents the local equilibrium behavior of RGG3.

Importantly, the use of all-atom MD simulations allows direct and accurate assessment of the configurational entropy change for RGG3 upon formation of the dense phase. Here, the decrease in the configurational entropy is almost two times smaller than in another recently studied IDR - Lge1 (1-80) - which undergoes robust LLPS *in vitro*^32^. Interestingly, the entropic penalty obtained for RGG3 is at the same time of a similar magnitude as that for Lge1 (1-80) Y2A mutant with impaired LLPS^32^. The latter suggests that IDRs showing robust LLPS may exhibit a certain threshold entropic penalty that facilitates condensate formation. An interesting question here concerns how the loss of configurational entropy in the condensed phase is thermodynamically compensated. If the protein configurational entropy decreases prominently upon condensation, it should at the same time increase for some other parts of the system. Specifically, the entropic penalty for RGG3 is linearly proportional to the relative reduction of its SASA. Thus, it is reasonable to assume that entropy of the solvent increases proportionally to the reduction of protein configurational entropy and this effect can be a driving force of protein condensation in the case of IDRs. The hypothesis should be systematically investigated in future work.

Overall, our results present a detailed atomistic view of the structure, dynamics and interactions within a model biomolecular condensate, and demonstrate the power of MD simulations in such applications. We hope that our results will stimulate further work on linking the microscopic, physicochemical aspects of disordered protein-protein interactions and condensate formation with their biological roles.

## Acknowledgments

Production simulations as well as parts of the analyses were carried out on the Vienna Scientific Cluster. Funding was provided by the FWF grants P30550 and P30680-B21 to BZ.

---

**Figure S1.**
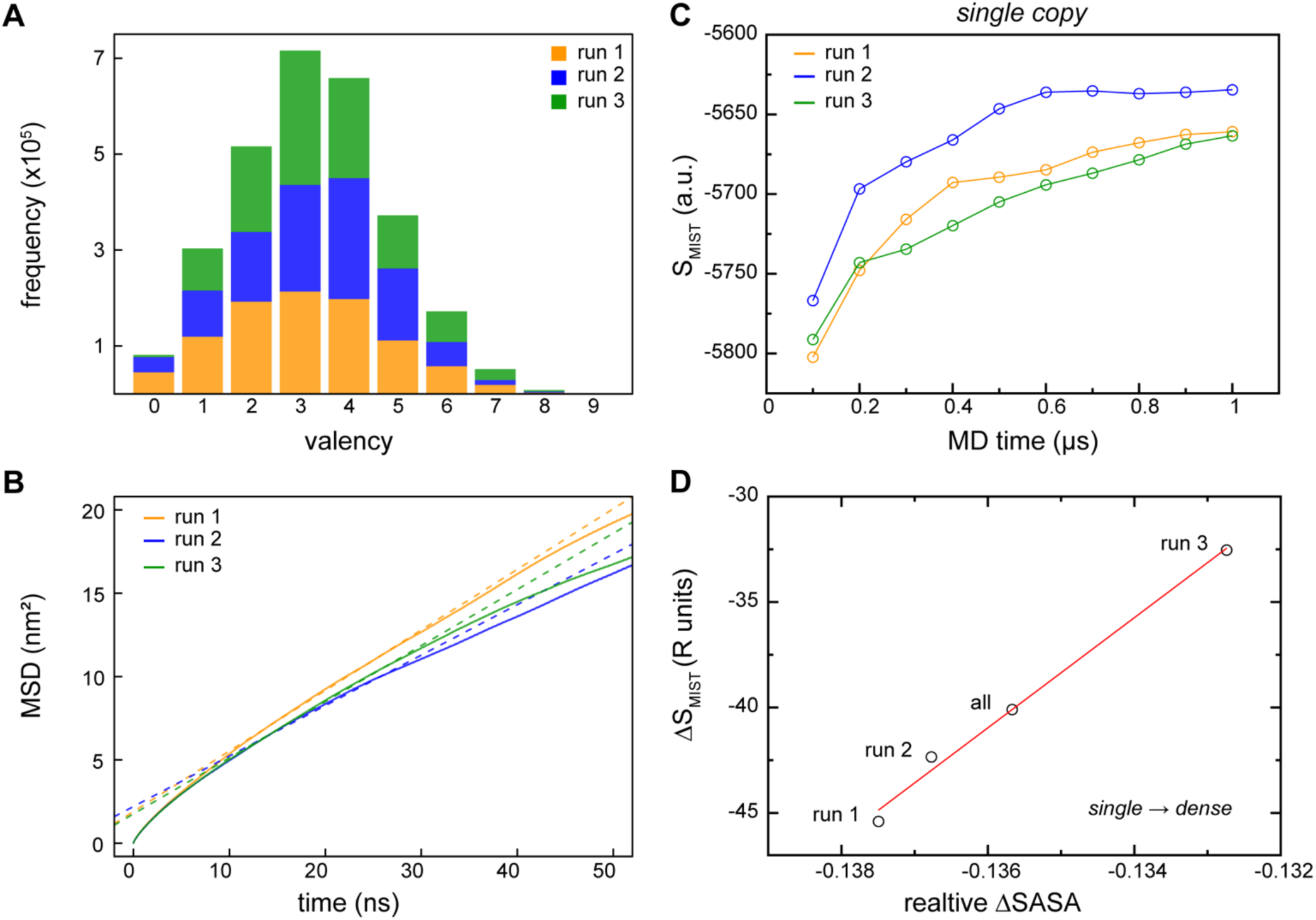
**A.** Distribution of average valencies over all 24 RGG3 copies at last 0.4 µs of each independent MD simulations of the dense phase. **B.** MSD-curve fitting for estimation of diffusion coefficients in the RGG3 dense phase with linear regime between 10-30 ns. MSD curves calculated using complete 1 µs MD trajectories. **C.** Convergence of RGG3 configurational entropy (*S_MIST_*) in the single-molecule context. Cumulative plots were generated using MIST approximation (see Methods) and a 100 ns time step. **D.** Linear regression between configurational entropy changes and corresponding changes in SASA of RGG3 upon transitioning into the dense phase. Relative entropy and SASA values were obtained using complete 1 µs MD trajectories.

**Figure S2.**
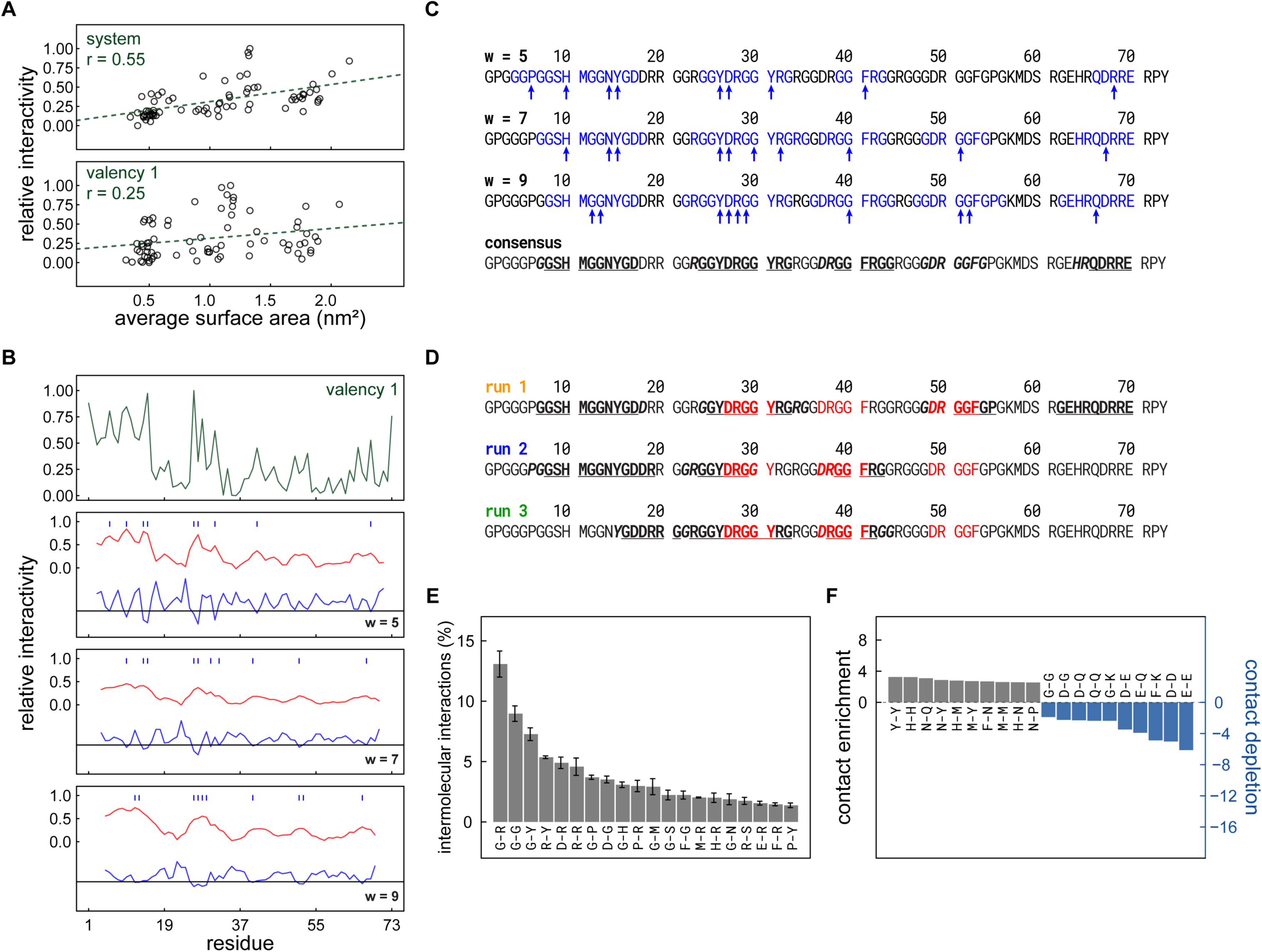
**A.** Relative interactivity at all valencies and valency 1 of different residues as a function of their average surface area at last 0.4 µs of MD simulations of the dense phase with the corresponding Pearson’s R. **B.** Visual example of peak picking using the Savitzky-Golay filter: The interactivity profile (top) is smoothened (red) and the second derivative (blue) is numerically determined using different window size, as indicated. The black lines indicate the distance of one standard deviation from the arithmetic mean of the second numerical derivatives, which is used as a cutoff for peak-picking. Blue bars above the smoothened interactivity profiles (red) indicate identified interactivity peaks. **C.** Sequence representation of Savitzky-Golay-identified interactivity peaks at different window sizes; arrows indicate the positions identified as peaks in the profile, while colored regions include residues that contributed to the peak signal. The bottom sequence shows the identified stickers in bold and underlined or bold and italic for full or reduced confidence, respectively. **D.** Comparison of stickers between replica simulations. **E.** The twenty most frequently occurring types of pairwise intermolecular contacts and the associated frequencies in percent. Error bars indicate standard deviation between the replicate trajectories. **F.** The ten most enriched and depleted interactions over all valencies.

**Figure S3.**
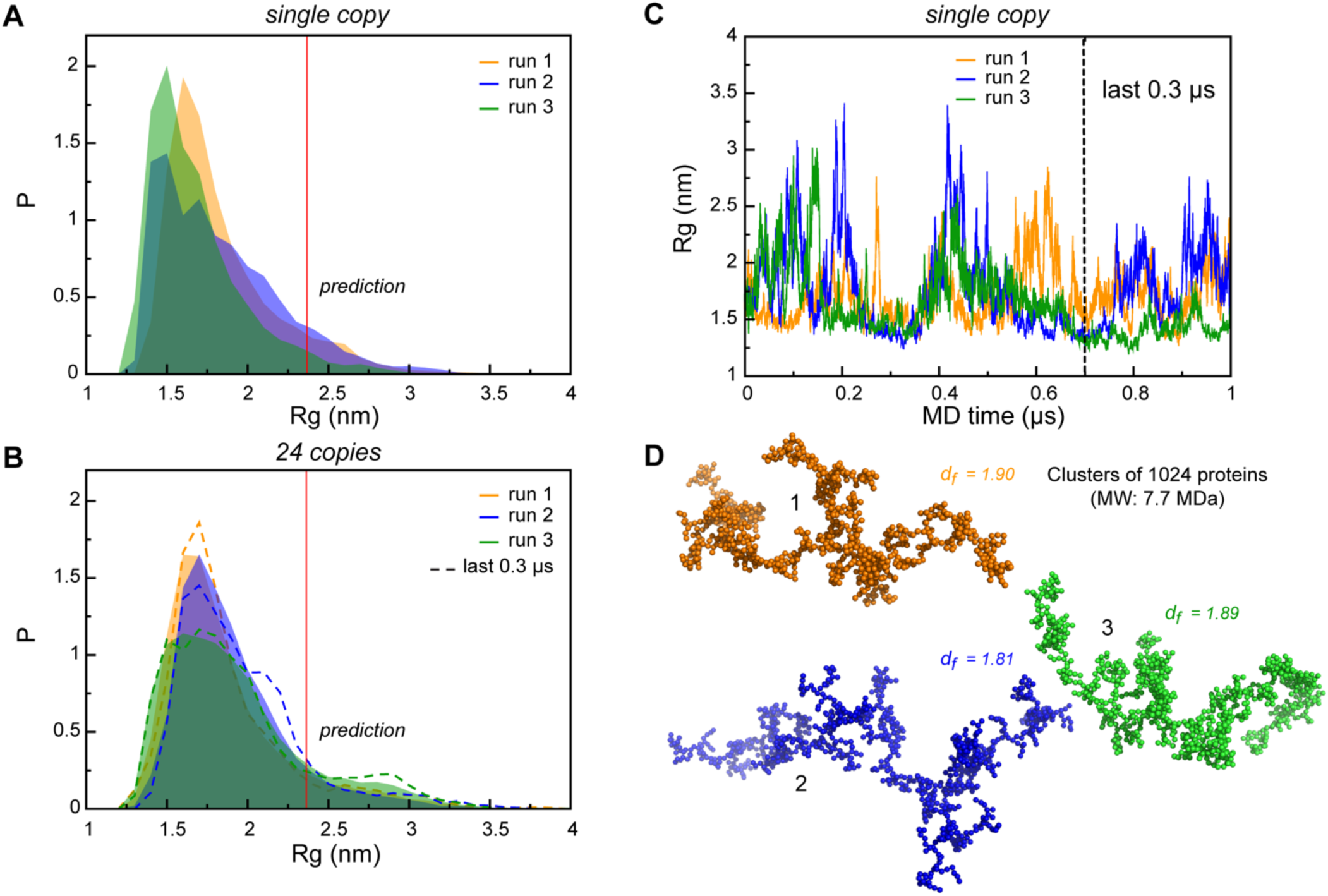
Distributions of radii of gyration (*Rg*) RGG3 in (**A**) single-copy and (**B**) multi-copy systems. Complete MD trajectories were used to collect *Rg* statistics. *Rg* distributions in the dense phase at last 0.3 µs of MD trajectories are shown with dashed lines. Theoretical *Rg_rc_* value for an 72-aa disordered protein chain (see Methods) is shown with a vertical red dashed line. **C.** Time evolution of *Rg* in RGG3 single copy MD simulations. **D.** Representative coarse-grained 1024 particle clusters obtained by FracVAL algorithm (see Methods). The clusters were reconstructed using the *d_F_* and the averaged *Rg* values at the last 0.3 µs of each independent MD simulations of the dense phase. Fractal dimensions (*d_F_*) corresponding to each cluster are indicated.

**Table S1.**
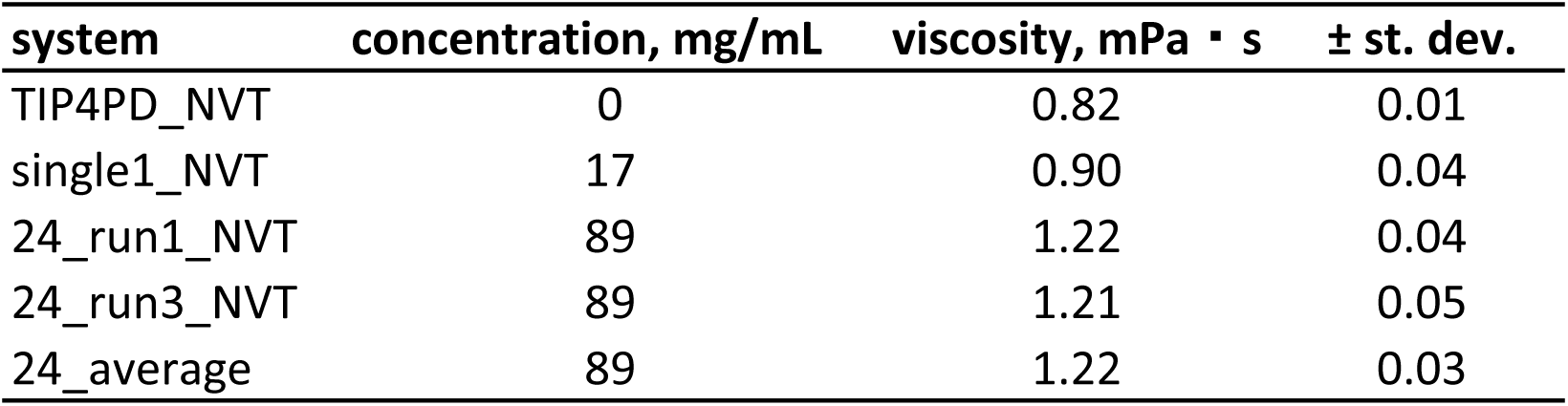
Shear viscosity values obtained from the analysis of different MD systems.

| system | concentration, mg/mL | viscosity, mPa · s | ± st. dev. |
| --- | --- | --- | --- |
| TIP4PD_NVT | 0 | 0.82 | 0.01 |
| single1_NVT | 17 | 0.90 | 0.04 |
| 24_run1_NVT | 89 | 1.22 | 0.04 |
| 24_run3_NVT | 89 | 1.21 | 0.05 |
| 24_average | 89 | 1.22 | 0.03 |

**Table S2.** Hotspot (sticker-sticker) interaction statistics.

| **description** | **fraction of all contacts** |
| --- | --- |
| Contacts involving stickers | 52.3% |
| Contacts between stickers | 43.2% |
| Contacts involving aromatic residues | 26.9% |
| Contacts between aromatic residues | 1.7% |
| Contacts involving arginines | 41.8% |
| Contacts between arginines and aromatics | 7.0% |

## References

1 Wright, P. E. & Dyson, H. J. Intrinsically disordered proteins in cellular signalling and regulation. Nature Reviews Molecular Cell Biology 16, 18–29, doi:10.1038/nrm3920 (2015).

2 Babu, M. M. The contribution of intrinsically disordered regions to protein function, cellular complexity, and human disease. Biochem Soc Trans 44, 1185–1200, doi:10.1042/bst20160172 (2016).

3 Tsang, B., Pritišanac, I., Scherer, S. W., Moses, A. M. & Forman-Kay, J. D. Phase Separation as a Missing Mechanism for Interpretation of Disease Mutations. Cell 183, 1742–1756, doi:10.1016/j.cell.2020.11.050 (2020).

4 McAffee, D. B. et al. Discrete LAT condensates encode antigen information from single pMHC:TCR binding events. Nature Communications 13, 7446, doi:10.1038/s41467-022-35093-9 (2022).

5 Bondos, S. E., Dunker, A. K. & Uversky, V. N. Intrinsically disordered proteins play diverse roles in cell signaling. Cell Communication and Signaling 20, 20, doi:10.1186/s12964-022-00821-7 (2022).

6 Banani, S. F., Lee, H. O., Hyman, A. A. & Rosen, M. K. Biomolecular condensates: organizers of cellular biochemistry. Nature Reviews Molecular Cell Biology 18, 285–298, doi:10.1038/nrm.2017.7 (2017).

7 Boeynaems, S. et al. Phase Separation in Biology and Disease; Current Perspectives and Open Questions. Journal of Molecular Biology 435, 167971, 10.1016/j.jmb.2023.167971 (2023).

8 Pappu, R. V., Cohen, S. R., Dar, F., Farag, M. & Kar, M. Phase Transitions of Associative Biomacromolecules. Chemical Reviews 123, 8945–8987, doi:10.1021/acs.chemrev.2c00814 (2023).

9 Brangwynne, C. P. et al. Germline P Granules Are Liquid Droplets That Localize by Controlled Dissolution/Condensation. Science 324, 1729–1732, doi:10.1126/science.1172046 (2009).

10 Molliex, A. et al. Phase Separation by Low Complexity Domains Promotes Stress Granule Assembly and Drives Pathological Fibrillization. Cell 163, 123–133, doi:10.1016/j.cell.2015.09.015 (2015).

11 Lafontaine, D. L. J., Riback, J. A., Bascetin, R. & Brangwynne, C. P. The nucleolus as a multiphase liquid condensate. Nature Reviews Molecular Cell Biology 22, 165–182, doi:10.1038/s41580-020-0272-6 (2021).

12 Zhu, L. & Brangwynne, C. P. Nuclear bodies: the emerging biophysics of nucleoplasmic phases. Current Opinion in Cell Biology 34, 23–30, 10.1016/j.ceb.2015.04.003 (2015).

13 Mitrea, D. M. & Kriwacki, R. W. Phase separation in biology; functional organization of a higher order. Cell Communication and Signaling 14, 1–1, doi:10.1186/s12964-015-0125-7 (2016).

14 Lyon, A. S., Peeples, W. B. & Rosen, M. K. A framework for understanding the functions of biomolecular condensates across scales. Nature Reviews Molecular Cell Biology 22, 215–235, doi:10.1038/s41580-020-00303-z (2021).

15 Uversky, V. N. Recent Developments in the Field of Intrinsically Disordered Proteins: Intrinsic Disorder-Based Emergence in Cellular Biology in Light of the Physiological and Pathological Liquid-Liquid Phase Transitions. Annu Rev Biophys 50, 135–156, doi:10.1146/annurev-biophys-062920-063704 (2021).

16 Alberti, S. & Hyman, A. A. Biomolecular condensates at the nexus of cellular stress, protein aggregation disease and ageing. Nature Reviews Molecular Cell Biology 22, 196–213, doi:10.1038/s41580-020-00326-6 (2021).

17 Holehouse, A. S. & Kragelund, B. B. The molecular basis for cellular function of intrinsically disordered protein regions. Nature Reviews Molecular Cell Biology 25, 187–211, doi:10.1038/s41580-023-00673-0 (2024).

18 Nomura, T. et al. Intranuclear aggregation of mutant FUS/TLS as a molecular pathomechanism of amyotrophic lateral sclerosis. J Biol Chem 289, 1192–1202, doi:10.1074/jbc.M113.516492 (2014).

19 Mészáros, B., Erdős, G. & Dosztányi, Z. IUPred2A: context-dependent prediction of protein disorder as a function of redox state and protein binding. Nucleic Acids Research 46, W329–W337, doi:10.1093/nar/gky384 (2018).

20 Emenecker, R. J., Griffith, D. & Holehouse, A. S. Metapredict: a fast, accurate, and easy-to-use predictor of consensus disorder and structure. Biophysical Journal 120, 4312–4319, 10.1016/j.bpj.2021.08.039 (2021).

21 Bugge, K. et al. Interactions by Disorder – A Matter of Context. Frontiers in Molecular Biosciences 7, doi:10.3389/fmolb.2020.00110 (2020).

22 Fuxreiter, M. Classifying the Binding Modes of Disordered Proteins. International Journal of Molecular Sciences 21, 8615–8615, doi:10.3390/ijms21228615 (2020).

23 Jahn, L. R., Marquet, C., Heinzinger, M. & Rost, B. Protein embeddings predict binding residues in disordered regions. Scientific Reports 14, 13566, doi:10.1038/s41598-024-64211-4 (2024).

24 Darling, A. L. & Uversky, V. N. Intrinsic Disorder and Posttranslational Modifications: The Darker Side of the Biological Dark Matter. Frontiers in Genetics 9, doi:10.3389/fgene.2018.00158 (2018).

25 Wang, J. et al. A Molecular Grammar Governing the Driving Forces for Phase Separation of Prion-like RNA Binding Proteins. Cell 174, 688–699.e616, doi:10.1016/j.cell.2018.06.006 (2018).

26 Vernon, R. M. et al. Pi-Pi contacts are an overlooked protein feature relevant to phase separation. eLife 7, doi:10.7554/eLife.31486 (2018).

27 Saar, K. L. et al. Learning the molecular grammar of protein condensates from sequence determinants and embeddings. Proceedings of the National Academy of Sciences 118, e2019053118, doi:10.1073/pnas.2019053118 (2021).

28 Flock, T., Weatheritt, R. J., Latysheva, N. S. & Babu, M. M. Controlling entropy to tune the functions of intrinsically disordered regions. Current Opinion in Structural Biology 26, 62–72, 10.1016/j.sbi.2014.05.007 (2014).

29 Soranno, A. et al. Integrated view of internal friction in unfolded proteins from single-molecule FRET, contact quenching, theory, and simulations. Proceedings of the National Academy of Sciences 114, E1833–E1839, doi:10.1073/pnas.1616672114 (2017).

30 Fleck, M., Polyansky, A. A. & Zagrovic, B. PARENT: A Parallel Software Suite for the Calculation of Configurational Entropy in Biomolecular Systems. J Chem Theory Comput 12, 2055–2065, doi:10.1021/acs.jctc.5b01217 (2016).

31 Fleck, M., Polyansky, A. A. & Zagrovic, B. Self-Consistent Framework Connecting Experimental Proxies of Protein Dynamics with Configurational Entropy. J Chem Theory Comput 14, 3796–3810, doi:10.1021/acs.jctc.8b00100 (2018).

32 Polyansky, A. A., Gallego, L. D., Efremov, R. G., Köhler, A. & Zagrovic, B. Protein compactness and interaction valency define the architecture of a biomolecular condensate across scales. eLife 12, e80038, doi:10.7554/eLife.80038 (2023).

33 Moses, D., Ginell, G. M., Holehouse, A. S. & Sukenik, S. Intrinsically disordered regions are poised to act as sensors of cellular chemistry. Trends in Biochemical Sciences 48, 1019–1034, 10.1016/j.tibs.2023.08.001 (2023).

34 Ghosh, C., Nagpal, S. & Muñoz, V. Molecular simulations integrated with experiments for probing the interaction dynamics and binding mechanisms of intrinsically disordered proteins. Current Opinion in Structural Biology 84, 102756, 10.1016/j.sbi.2023.102756 (2024).

35 Piana, S., Donchev, A. G., Robustelli, P. & Shaw, D. E. Water Dispersion Interactions Strongly Influence Simulated Structural Properties of Disordered Protein States. The Journal of Physical Chemistry B 119, 5113–5123, doi:10.1021/jp508971m (2015).

36 Piana, S., Robustelli, P., Tan, D., Chen, S. & Shaw, D. E. Development of a Force Field for the Simulation of Single-Chain Proteins and Protein–Protein Complexes. Journal of Chemical Theory and Computation 16, 2494–2507, doi:10.1021/acs.jctc.9b00251 (2020).

37 Gopal, S. M. et al. Conformational Preferences of an Intrinsically Disordered Protein Domain: A Case Study for Modern Force Fields. J Phys Chem B 125, 24–35, doi:10.1021/acs.jpcb.0c08702 (2021).

38 Wang, W. Recent advances in atomic molecular dynamics simulation of intrinsically disordered proteins. Physical Chemistry Chemical Physics 23, 777–784, doi:10.1039/D0CP05818A (2021).

39 Kasahara, K., Terazawa, H., Takahashi, T. & Higo, J. Studies on Molecular Dynamics of Intrinsically Disordered Proteins and Their Fuzzy Complexes: A Mini-Review. Comput Struct Biotechnol J 17, 712–720, doi:10.1016/j.csbj.2019.06.009 (2019).

40 Bastida, A., Zúñiga, J., Fogolari, F. & Soler, M. A. Statistical accuracy of molecular dynamics-based methods for sampling conformational ensembles of disordered proteins. Physical Chemistry Chemical Physics, doi:10.1039/D4CP02564D (2024).

41 Zhu, J., Salvatella, X. & Robustelli, P. Small molecules targeting the disordered transactivation domain of the androgen receptor induce the formation of collapsed helical states. Nature Communications 13, 6390, doi:10.1038/s41467-022-34077-z (2022).

42 Galvanetto, N. et al. Extreme dynamics in a biomolecular condensate. Nature 619, 876–883, doi:10.1038/s41586-023-06329-5 (2023).

43 Rauscher, S. & Pomès, R. The liquid structure of elastin. eLife 6, e26526, doi:10.7554/eLife.26526 (2017).

44 Paloni, M., Bailly, R., Ciandrini, L. & Barducci, A. Unraveling Molecular Interactions in Liquid-Liquid Phase Separation of Disordered Proteins by Atomistic Simulations. J Phys Chem B 124, 9009–9016, doi:10.1021/acs.jpcb.0c06288 (2020).

45 Flores-Solis, D. et al. Driving forces behind phase separation of the carboxy-terminal domain of RNA polymerase II. Nature Communications 14, 5979, doi:10.1038/s41467-023-41633-8 (2023).

46 Dignon, G. L., Zheng, W., Best, R. B., Kim, Y. C. & Mittal, J. Relation between single-molecule properties and phase behavior of intrinsically disordered proteins. Proceedings of the National Academy of Sciences 115, 9929–9934, doi:10.1073/pnas.1804177115 (2018).

47 Chou, H. Y. & Aksimentiev, A. Single-Protein Collapse Determines Phase Equilibria of a Biological Condensate. J Phys Chem Lett 11, 4923–4929, doi:10.1021/acs.jpclett.0c01222 (2020).

48 Bremer, A. et al. Deciphering how naturally occurring sequence features impact the phase behaviours of disordered prion-like domains. Nature Chemistry 14, 196–207, doi:10.1038/s41557-021-00840-w (2022).

49 Lotthammer, J. M., Ginell, G. M., Griffith, D., Emenecker, R. J. & Holehouse, A. S. Direct prediction of intrinsically disordered protein conformational properties from sequence. Nature Methods 21, 465–476, doi:10.1038/s41592-023-02159-5 (2024).

50 Milles, S. & Lemke, E. A. Mapping Multivalency and Differential Affinities within Large Intrinsically Disordered Protein Complexes with Segmental Motion Analysis. Angewandte Chemie International Edition 53, 7364–7367, 10.1002/anie.201403694 (2014).

51 Fung, H. Y. J., Birol, M. & Rhoades, E. IDPs in macromolecular complexes: the roles of multivalent interactions in diverse assemblies. Curr Opin Struct Biol 49, 36–43, doi:10.1016/j.sbi.2017.12.007 (2018).

52 Martin, E. W. et al. Valence and patterning of aromatic residues determine the phase behavior of prion-like domains. Science 367, 694–699, doi:10.1126/science.aaw8653 (2020).

53 Sipko, E. L., Chappell, G. F. & Berlow, R. B. Multivalency emerges as a common feature of intrinsically disordered protein interactions. Current Opinion in Structural Biology 84, 102742, 10.1016/j.sbi.2023.102742 (2024).

54 De La Cruz, N. et al. Disorder-mediated interactions target proteins to specific condensates. Molecular Cell, 10.1016/j.molcel.2024.08.017 (2024).

55 Saar, K. L. et al. Protein Condensate Atlas from predictive models of heteromolecular condensate composition. Nature Communications 15, 5418, doi:10.1038/s41467-024-48496-7 (2024).

56 Alshareedah, I. et al. Sequence-specific interactions determine viscoelasticity and ageing dynamics of protein condensates. Nature Physics, doi:10.1038/s41567-024-02558-1 (2024).

57 Sundaravadivelu Devarajan, D., et al. Sequence-dependent material properties of biomolecular condensates and their relation to dilute phase conformations. Nature Communications 15, 1912, doi:10.1038/s41467-024-46223-w (2024).

58 Kozak, F. et al. An Atomistic View on the Mechanism of Diatom Peptide-Guided Biomimetic Silica Formation. Advanced Science n/a, 2401239, 10.1002/advs.202401239 (2024).

59 Mittag, T. & Pappu, R. V. A conceptual framework for understanding phase separation and addressing open questions and challenges. Mol Cell 82, 2201–2214, doi:10.1016/j.molcel.2022.05.018 (2022).

60 Kar, M. et al. Solutes unmask differences in clustering versus phase separation of FET proteins. Nature Communications 15, 4408, doi:10.1038/s41467-024-48775-3 (2024).

61 Gil-Garcia, M. et al. Local environment in biomolecular condensates modulates enzymatic activity across length scales. Nature Communications 15, 3322, doi:10.1038/s41467-024-47435-w (2024).

62 Crozat, A., Åman, P., Mandahl, N. & Ron, D. Fusion of CHOP to a novel RNA-binding protein in human myxoid liposarcoma. Nature 363, 640–644, doi:10.1038/363640a0 (1993).

63 Tan, A. Y., Riley, T. R., Coady, T., Bussemaker, H. J. & Manley, J. L. TLS/FUS (translocated in liposarcoma/fused in sarcoma) regulates target gene transcription via single-stranded DNA response elements. Proceedings of the National Academy of Sciences of the United States of America 109, 6030–6035, doi:10.1073/pnas.1203028109 (2012).

64 Ederle, H. & Dormann, D. TDP-43 and FUS en route from the nucleus to the cytoplasm. FEBS letters 591, 1489–1507, doi:10.1002/1873-3468.12646 (2017).

65 Schoen, M. et al. Super-Resolution Microscopy Reveals Presynaptic Localization of the ALS/FTD Related Protein FUS in Hippocampal Neurons. Frontiers in cellular neuroscience 9, 496–496, doi:10.3389/fncel.2015.00496 (2015).

66 Patel, A. et al. A Liquid-to-Solid Phase Transition of the ALS Protein FUS Accelerated by Disease Mutation. Cell 162, 1066–1077, doi:10.1016/j.cell.2015.07.047 (2015).

67 Deng, Q. et al. FUS is Phosphorylated by DNA-PK and Accumulates in the Cytoplasm after DNA Damage. Journal of Neuroscience 34, 7802–7813, doi:10.1523/JNEUROSCI.0172-14.2014 (2014).

68 Banani, S. F. et al. Compositional Control of Phase-Separated Cellular Bodies. Cell 166, 651–663, doi:10.1016/j.cell.2016.06.010 (2016).

69 Yoshizawa, T. et al. Nuclear Import Receptor Inhibits Phase Separation of FUS through Binding to Multiple Sites. Cell 173, 693–705.e622, doi:10.1016/j.cell.2018.03.003 (2018).

70 Qamar, S. et al. FUS Phase Separation Is Modulated by a Molecular Chaperone and Methylation of Arginine Cation-π Interactions. Cell 173, 720–734.e715, doi:10.1016/j.cell.2018.03.056 (2018).

71 Schuster, B. S. et al. Identifying sequence perturbations to an intrinsically disordered protein that determine its phase-separation behavior. Proceedings of the National Academy of Sciences 117, 11421–11431, doi:10.1073/pnas.2000223117 (2020).

72 Zheng, W. et al. Molecular Details of Protein Condensates Probed by Microsecond Long Atomistic Simulations. J Phys Chem B 124, 11671–11679, doi:10.1021/acs.jpcb.0c10489 (2020).

73 Hong, Y. et al. Hydrophobicity of arginine leads to reentrant liquid-liquid phase separation behaviors of arginine-rich proteins. Nature Communications 13, 7326, doi:10.1038/s41467-022-35001-1 (2022).

74 Hofweber, M. et al. Phase Separation of FUS Is Suppressed by Its Nuclear Import Receptor and Arginine Methylation. Cell 173, 706–719.e713, doi:10.1016/j.cell.2018.03.004 (2018).

75 Baade, I. et al. The RNA-binding protein FUS is chaperoned and imported into the nucleus by a network of import receptors. Journal of Biological Chemistry 296, 100659–100659, doi:10.1016/j.jbc.2021.100659 (2021).

76 Van Der Spoel, D. et al. GROMACS: Fast, flexible, and free. Journal of Computational Chemistry 26, 1701–1718, doi:10.1002/jcc.20291 (2005).

77 GROMACS User Manual version 5.1.4 (2016).

78 Lindorff-Larsen, K. et al. Improved side-chain torsion potentials for the Amber ff99SB protein force field. Proteins: Structure, Function, and Bioinformatics 78, 1950–1958, doi:10.1002/prot.22711 (2010).

79 RSutdio: Integrated Development for R. RStudio, Inc (RStudio, Inc., Boston, MA, 2016).

80 Wickham, H. ggplot2. (Springer International Publishing, 2016).

81 Jaki, T. & Wolfsegger, M. J. Estimation of pharmacokinetic parameters with the R package PK. Pharmaceutical Statistics 10, 284–288, doi:10.1002/pst.449 (2011).

82 Källberg, M. et al. Template-based protein structure modeling using the RaptorX web server. Nature Protocols 7, 1511–1522, doi:10.1038/nprot.2012.085 (2012).

83 Kelley, L. A., Mezulis, S., Yates, C. M., Wass, M. N. & Sternberg, M. J. E. The Phyre2 web portal for protein modeling, prediction and analysis. Nature Protocols 10, 845–858, doi:10.1038/nprot.2015.053 (2015).

84 Hess, B., Bekker, H., Berendsen, H. J. C. & Fraaije, J. G. E. M. LINCS: A linear constraint solver for molecular simulations. Journal of Computational Chemistry 18, 1463–1472, doi:10.1002/(SICI)1096-987X(199709)18:12<1463::AID-JCC4>3.0.CO;2-H (1997).

85 Hoover, W. G. Canonical dynamics: Equilibrium phase-space distributions. Physical Review A 31, 1695–1697, doi:10.1103/PhysRevA.31.1695 (1985).

86 Berendsen, H. J. C., Postma, J. P. M., van Gunsteren, W. F., DiNola, A. & Haak, J. R. Molecular dynamics with coupling to an external bath. The Journal of Chemical Physics 81, 3684–3690, doi:10.1063/1.448118 (1984).

87 Parrinello, M. & Rahman, A. Polymorphic transitions in single crystals: A new molecular dynamics method. Journal of Applied Physics 52, 7182–7190, doi:10.1063/1.328693 (1981).

88 von Bülow, S., Siggel, M., Linke, M. & Hummer, G. Dynamic cluster formation determines viscosity and diffusion in dense protein solutions. Proceedings of the National Academy of Sciences 116, 9843–9852, doi:10.1073/pnas.1817564116 (2019).

89 Hess, B. Determining the shear viscosity of model liquids from molecular dynamics simulations. The Journal of Chemical Physics 116, 209–217, doi:10.1063/1.1421362 (2002).

90 Yeh, I.-C. & Hummer, G. System-Size Dependence of Diffusion Coefficients and Viscosities from Molecular Dynamics Simulations with Periodic Boundary Conditions. The Journal of Physical Chemistry B 108, 15873–15879, doi:10.1021/jp0477147 (2004).

91 Fennell, C. J., Ghousifam, N., Haseleu, J. M. & Gappa-Fahlenkamp, H. Computational Signaling Protein Dynamics and Geometric Mass Relations in Biomolecular Diffusion. The Journal of Physical Chemistry B 122, 5599–5609, doi:10.1021/acs.jpcb.7b11846 (2018).

92 Lohninger, H.

93 Taylor, J. R. An introduction to error analysis : the study of uncertainties in physical measurements. 2. ed. edn, (Univ. Science Books, 1997).

94 King, B. M., Silver, N. W. & Tidor, B. Efficient calculation of molecular configurational entropies using an information theoretic approximation. J Phys Chem B 116, 2891–2904, doi:10.1021/jp2068123 (2012).

95 Morán, J., Fuentes, A., Liu, F. & Yon, J. FracVAL: An improved tunable algorithm of cluster–cluster aggregation for generation of fractal structures formed by polydisperse primary particles. Computer Physics Communications 239, 225–237, 10.1016/j.cpc.2019.01.015 (2019).

96 Kang, J., Lim, L. & Song, J. ATP enhances at low concentrations but dissolves at high concentrations liquid-liquid phase separation (LLPS) of ALS/FTD-causing FUS. Biochemical and Biophysical Research Communications 504, 545–551, doi:10.1016/j.bbrc.2018.09.014 (2018).

97 Ruff, K. M., Roberts, S., Chilkoti, A. & Pappu, R. V. Advances in Understanding Stimulus-Responsive Phase Behavior of Intrinsically Disordered Protein Polymers. Journal of molecular biology 430, 4619–4635, doi:10.1016/j.jmb.2018.06.031 (2018).

98 Monti, M. et al. Accurate Predictions of Liquid-Liquid Phase Separating Proteins at Single Amino Acid Resolution. bioRxiv, 2024.2007.2019.602785, doi:10.1101/2024.07.19.602785 (2024).

99 Kumar, K. et al. Cation–π interactions in protein–ligand binding: theory and data-mining reveal different roles for lysine and arginine. Chemical Science 9, 2655–2665, doi:10.1039/C7SC04905F (2018).

100 Fuxreiter, M. & Tompa, P. in Fuzziness: Structural Disorder in Protein Complexes (eds Monika Fuxreiter & Peter Tompa) 1–14 (Springer US, 2012).

101 Bernadó, P. & Blackledge, M. A Self-Consistent Description of the Conformational Behavior of Chemically Denatured Proteins from NMR and Small Angle Scattering. Biophysical Journal 97, 2839–2845, doi:10.1016/j.bpj.2009.08.044 (2009).

102 Kar, M. et al. Phase-separating RNA-binding proteins form heterogeneous distributions of clusters in subsaturated solutions. Proceedings of the National Academy of Sciences 119, e2202222119, doi:doi:10.1073/pnas.2202222119 (2022).

